# Comprehensive Pan-Cancer Analysis Reveals the Prognostic, Molecular, and Immunological Significance of EPS8

**DOI:** 10.64898/2026.09.13.751329

**Authors:** Adiba Juoairia, Ayesha Akter, Rawaz Jahan Nima, Md Azizul Haque

**Affiliations:** Computer Science and Engineering, Manarat International University, Dhaka, Bangladesh; Computer Science and Engineering, BRAC University, Dhaka, Bangladesh; Department of Forensic Medicine and Toxicology, Ibrahim Medical College, Dhaka, Bangladesh; Department of Biotechnology, Yeungnam University, Gyeongsan-si, South Korea

**Keywords:** Cancer biology, Human tumors, EPS8, Biomarker, Pan-cancer analysis, Oncogenesis

## Abstract

**Background:** Epidermal Growth Factor Receptor Pathway Substrate 8 (EPS8) is believed to function as a tumor driver; however, to understand its molecular characteristics across tumors, a comprehensive pan-cancer analysis of EPS8 is lacking.

**Objective:** This study aimed to investigate the prognostic, molecular, and immunological significance of EPS8 across multiple human cancers.

**Methods:** Comprehensive analyses were conducted using GEPIA2, UALCAN, TIMER2.0, cBioPortal, SMART, GSCA, Enrichr, and the TCGAplot R package to evaluate survival prognosis, gene expression, DNA methylation, immune infiltration, proteomic expression, tumor mutational burden (TMB), microsatellite instability (MSI), genetic alterations, drug sensitivity, and enriched pathways.

**Results:** EPS8 overexpression in LGG (*p* = 7.2 × 10^−5^) and PAAD (*p* = 9.4 × 10^−6^) was significantly associated with poor overall survival and disease-free survival. Amplification was the most common genetic alteration, with alteration frequencies approaching 6% in TGCT and UCEC. In KIRC and PAAD, EPS8 expression correlated with tumor stage. Immune infiltration analysis revealed significant associations between EPS8 expression and immune cells. EPS8 expression also showed significant correlations with both TMB and MSI in STAD and ESCA. Enrichment analysis indicated an association between EPS8 and regulation of the actin cytoskeleton and similar pathways.

**Conclusion:** These findings suggest that EPS8 has prognostic and immunological significance across multiple cancer types and may serve as a potential biomarker. The observed associations with immune-related features and drug response provide a basis for future experimental studies to evaluate its role in cancer biology and its potential relevance to immunotherapy.

## 1 Introduction

The cancer problem is a leading cause of death globally. In 2020, 19.3 million new cancer cases were diagnosed, and nearly 10 million cancer deaths were recorded globally. By 2040, an additional 47% increase in new cancer cases is projected, totaling 28.4 million [1]. Breast cancer is diagnosed the most among all cancers; however, lung cancer is the major cause of death, followed by colorectal cancer, liver cancer, and stomach cancer [2]. This is becoming a shifting problem in Asia, as the cancer burden continues to increase. A study reports that over 50% of cancer deaths and almost half of all new cancer cases globally occur in this region [3]. Specifically in China, the incidence and mortality rates from cancer are increasing, especially due to lung and colorectal cancers, which are the most common [4]. This reflects the regional burden imposed by large populations, diverse cancer risk factors, and limited access to prevention and early detection measures [5]. Even with advances in cancer treatment, challenges remain due to inadequate cancer data, low screening rates, and limited economic accessibility to modern cancer treatment in low-and middle-income countries [6]. To address the rising cancer burden, research limited to conventional clinical data is no longer sufficient, motivating the need for in-depth cancer research to determine cancer-promoting mechanisms at the genetic and epigenetic levels.

Of the many molecular systems, epidermal growth factor receptor kinase substrate 8 (EPS8) is a member of a specialized class of scaffold proteins that includes EPS8, EPS8L1, EPS8L2, and EPS8L3 [7]. These proteins contain a modular structure consisting of one PH domain and one SH3 domain, and are part of the EGFR pathway [8]. They serve as signaling adaptors that connect growth factor receptor activation to remodeling of the actin cytoskeleton. EPS8 influences actin dynamics through two mechanisms: capping the barbed ends of actin filaments and bundling actin filaments, thereby influencing cell motility and morphogenesis [7]. At the cellular level, EPS8 also functions as a scaffold that integrates signaling pathways controlling cell proliferation, migration, and receptor trafficking [7]. Within the EPS8 family, the primary site of action is growth factor-initiated signaling, particularly the EGFR-Ras-Rac axis [7]. EPS8 enhances signaling pathways that promote malignant phenotypes by acting as a substrate of EGFR kinase, which can lead to repression of tumor suppressors [9]. EPS8 also contributes to DNA damage repair mechanisms responsible for chemoresistance in cancers such as NSCLC by modulating the ATM-p53 pathway [10]. EPS8L2 and EPS8L3, two other members of the EPS8 family, are involved in the progression of colorectal and hepatocellular carcinoma [11].

EPS8, the founding member of the EPS8 family, encodes a protein of approximately 97 kDa that serves as a substrate for receptor tyrosine kinases [7]. It is a multifunctional scaffolding protein that links EGFR signaling to downstream pathways regulating cellular mitosis, differentiation, and the malignant behaviors of tumor proliferation, invasion, metastasis, and drug resistance [7, 12]. EPS8 distinguishes itself from other EGFR pathway components as a unique oncoprotein that integrates signaling, receptor trafficking, and cytoskeletal remodeling across various tumors [12]. It can also act as a molecular switch that prolongs mitogenic signaling at the plasma membrane [9], while its interaction with E3b1/Sos1 drives Rac-mediated actin remodeling, establishing EPS8 as a novel actin-capping protein essential for tumor invasion and migration [9]. Furthermore, EPS8 possesses a non-canonical SH3 domain that recognizes unique PXXDY motifs, and has been identified as an independent predictor of poor overall survival (OS) in various cancers. Its significance is further supported by frequent overexpression in solid tumors and hematological malignancies even when EGFR levels remain constant [13].

EPS8 is consistently overexpressed across various cancers. In malignant melanoma, it activates the Hedgehog pathway through degradation of Ptch1, increasing tumor growth and worsening prognosis [14]. In prostate cancer, EPS8 overexpression is associated with enzalutamide resistance and acquisition of an epithelial-to-mesenchymal transition (EMT) phenotype [15]. In breast cancer, EPS8 overexpression is linked to heightened proliferation and migration, partial EMT-like features, and diminished chemotherapy response [12]. In esophageal cancer, EPS8 shows higher expression in tumor cells than in normal tissue, affecting proliferation and metastasis [16]. In oral squamous cell carcinoma, significant cytoplasmic staining was observed in 39% of tumors, with gene expression enhanced more than 5-fold in OSCC cell lines compared to normal keratinocytes [17]. In pancreatic cancer, EPS8 is a key marker promoting cell viability, clonogenicity, migration, and invasion [18]. In glioblastoma, EPS8 is a putative target that promotes tumor progression through the PI3K/Akt pathway [12, 19]. While these individual studies establish EPS8’s oncogenic role in specific tumor types, no study to date has examined EPS8 systematically across the cancer spectrum.

A pan-cancer analysis is the systematic evaluation of molecular features across multiple tumor types, providing a powerful framework to identify shared oncogenic mechanisms and context-specific effects that may be overlooked in single-cancer studies. This approach integrates genomic, transcriptomic, proteomic, and clinical data from large public repositories (e.g., TCGA, GTEx, CPTAC) to uncover tissue-selective mechanisms, novel pathway dysregulations, and clinically relevant biomarkers. Such studies have previously enabled the classification of universal biomarkers and therapeutic targets, such as the molecular chaperones CCT2 and CCT8, across multiple cancer types [19, 20], underscoring the value of this approach for characterizing a protein’s unified expression, alteration, and clinical relevance across malignancies.

Despite this established oncogenic role of EPS8 in individual cancers, no pan-cancer study has yet examined EPS8 comprehensively across tumor types, including its expression patterns, genomic alterations, epigenetic regulation, associations with the immune microenvironment, and prognostic value. This gap is significant because findings from single-cancer studies cannot establish whether EPS8 acts as a universal oncogenic driver or exerts tumor-specific effects, information that is essential before EPS8 can be pursued as a cross-cancer therapeutic target or biomarker. This study addresses that gap by performing the first pan-cancer characterization of EPS8, integrating multi-omics and clinical data from TCGA and GTEx to clarify its molecular characteristics, regulatory mechanisms, and clinical significance across tumor types.

Specifically, this study aims to: (1) characterize EPS8 mRNA and protein expression across tumor and normal tissues; (2) evaluate EPS8 genomic and epigenetic alterations, including mutation frequency, copy number variation, and promoter methylation; (3) assess associations between EPS8 expression and the tumor immune microenvironment, including immune cell infiltration and immune-related gene signatures; and (4) determine the prognostic value of EPS8 across cancer types using various analyses. Together, these objectives will clarify whether EPS8 functions as a pan-cancer oncogene or a tumor-specific factor, and establish its potential as a prognostic marker and therapeutic target to support future precision oncology strategies.

## 2 Method

### 2.1 Dataset Information and workflow

This study used The Cancer Genome Atlas TCGA, a public dataset providing genomic and epigenomic data across 33 cancer types, to conduct a comprehensive pan-cancer analysis of EPS8 [21]. The workflow, as demonstrated in Figure 1, included evaluation of EPS8 gene and protein expression, survival prognosis, DNA methylation, genetic alterations, immune infiltration, tumor mutational burden (TMB), microsatellite instability (MSI), drug response, single-cell expression, and pathway enrichment to comprehensively characterize the molecular, prognostic, and immunological significance of EPS8 across multiple cancer types.

**Figure 1:**
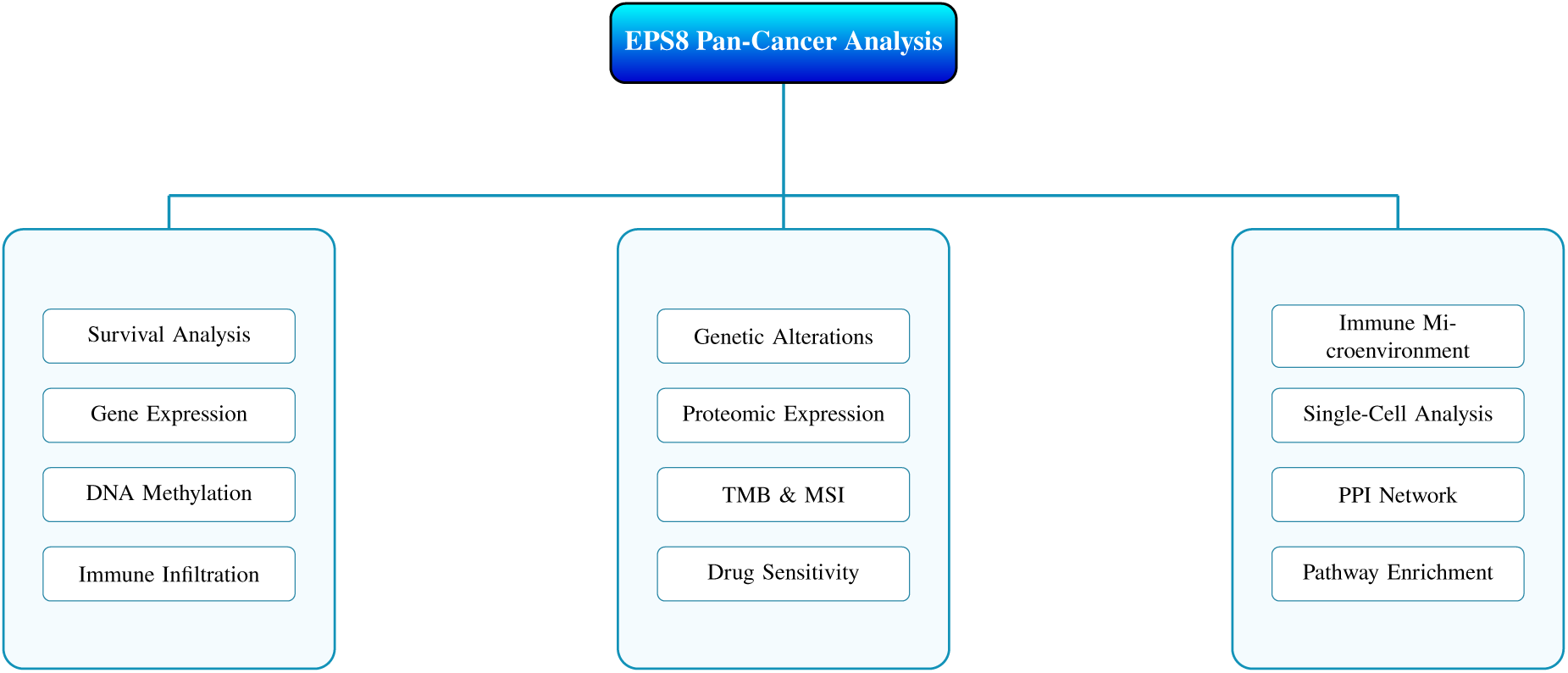
Overview of the semantic workflow for the EPS8 pan-cancer analysis.

### 2.2 Survival Prognosis Analysis

The “Survival analysis and survival map” module of the GEPIA2, a TCGA/GTEx-based expression profiling tool [22], was used for Overall Survival (OS) and Disease-Free Survival (DFS) plots, alongside a survival heatmap of EPS8 across TCGA. The expression threshold was set at 50%, with the group cut-off defined by the median. The survival analysis was done using both DFS and OS to find the prognostic value of EPS8 [23].

### 2.3 EPS8 expression analysis

In this study, we deployed a web server for gene expression analysis, TIMER 2.0. The “Gene DE” module was used to conduct expression analysis to systematically study the differences in EPS8 expression between tumor and normal tissues [24]. We also utilized the “Expression DIY-Box Plot” module of the GEPIA2 to validate the expression dependencies across the TCGA and GTEx databases with options set as P-value cut-off=0.01, log2FC cut-off=TRUE, Jitter size=0.4. We also explored EPS8 expression across the multiple clinical stages of cancer (stages I, II, III, and IV) from the TCGA database using the “Expression analysis-Stage Plot” module of GEPIA2. Additionally, EPS8 expression variation across various cancer types was also obtained using the UALCAN, an interactive web portal for analyzing Tumor subgroups [25].

### 2.4 DNA methylation analysis

Promoter methylation analysis for EPS8 in normal and cancer tissues was performed using the “TCGA analysis-methylation” module in the UALCAN database. Furthermore, a web-based tool for DNA methylation analysis and visualization, SMART, was used to create visualizations of the chromosomal distribution of the probes, aggregated methylation of CpG, and DNA methylation sites [26].

### 2.5 Immune infiltration analysis

To study immune infiltration associated with EPS8 across TCGA, the TIMER2.0 “immune-gene” module was used. Among the immune cells studied were cancer-associated fibroblasts (CAFs) and T follicular helper cells (Tfh). The level of immune infiltration was determined utilizing the EPIC, MCPCOUNTER, XCELL, TIDE, CIBERSORT, and CIBERSORT-ABS algorithms. We used a Purity-adjusted Spearman’s rank correlation test to get the p-value and the corresponding partial correlation coefficient (corr) [27].

### 2.6 Gene alteration analysis

The cBioPortal (An open-access web portal for analyzing multidimensional cancer genomics data) was used to obtain data related to genetic alterations of EPS8 to perform alteration analysis [28]. The “Cancer Types Summary” module provided the data on alteration frequency, mutation type, and CNA (Copy number alterations) for EPS8 in TCGA tumors. The mutated site for EPS8 was from the “Mutations” module. We used the “TCGA-pan cancer atlas” and “Query by gene” to obtain data regarding Overall Survival (OS), Disease-Free Survival (DFS), Progression-Free Survival (PFS), and Disease-Specific Survival (DSS) of the TCGA tumors with EPS8 genetic alterations compared to those without genetic alteration in the “Comparison/Survival” module. We also visualized log-rank P-values along with Kaplan-Meier plots.

### 2.7 Gene Proteomics analysis

The UALCAN database was deployed to understand the difference between protein expression in normal and tumor tissues. UALCAN allows proteomics analysis from the International Cancer Proteogenome Consortium (ICPC) and the Clinical Proteomic Tumor Analysis Consortium (CPTAC) datasets. UALCAN gives a protein expression analysis of 14 tumor types [29].

### 2.8 TMB and MSI correlation analysis of EPS8

Tumor mutation burden (TMB) is an emerging addition to the arsenal of predictive biomarkers for the immune checkpoint inhibitors (ICIs). When TMB is higher, there is a stronger neoantigen load, tumor immunogenicity, and increased probability of T cell recognition and response to ICIs [30]. Another example of a genetic alteration is microsatellite instability (MSI). During tumorigenesis, the DNA repair mechanism is disrupted, resulting in alterations in microsatellite sequence lengths, which result in instability of the microsatellite involved, known as MSI [31]. The TMB and MSI correlation with EPS8 expression analyses was done using the Pearson correlation method, and a visualization was demonstrated through a radar plot created using the TCGAplot R package, which is a package for visualization and analysis of TCGA gene expression and clinical data [32, 33].

### 2.9 Single-cell functional analysis

We deployed the “Gene Exploration” module in the TISCH2, a database for exploring TME using single-cell RNA-seq datasets, to perform the single-cell functional analysis [34] and configured the following parameters: gene name “EPS8”, cell type “Celltype (major-lineage)”, and cancer types including BRCA, BLCA, CHOL, ESCA, HNSC, KIRC, LIHC, OV, STAD, SKCM, PRAD, PAAD, UVM, THCA, and UCEC.

### 2.10 Assessment of immune microenvironment composition

Immune microenvironment assessment was done to understand the correlation between EPS8 and immune cells, immune checkpoints, immune score, and immune regulatory genes (immunoinhibitory, immunostimulatory). The TCGAplot was used to generate the heatmaps showcasing the correlation [32].

### 2.11 EPS8 related drug responsne analysis

To understand the correlation between drug response and EPS8 expression, we used the GSCA database [35]. It is a database that combines genomics data across TCGA cancer types, drug response information of over 750 small-molecule drugs from Genomics of Drug Sensitivity in Cancer (GDSC), and The Cancer Therapeutics Response Portal (CTRP) [36]. Drug sensitivity associations were considered significant at FDR ≤ 0.05. Correlation coefficients ranged from −0.4 to 0.4 for GDSC and from 0.0 to 0.5 for CTRP, with stronger positive correlations shown in red and stronger negative correlations shown in blue. Circle size represents statistical significance, with larger circles indicating higher significance (− log_10_(FDR) = 10).

### 2.12 Gene enrichment analysis

In this study, we deployed STRING, a database to visualize protein–protein interaction (PPI) networks, to investigate the PPI of EPS8. The parameter configuration was: the meaning of network edges: “confidence”, and all available sources were selected as active interaction sources; the confidence was set to low confidence, and the maximum number of interactors was 50 [37]. Next, we utilized the “Similar Gene Detection” from GEPIA2 to identify the top 100 co-expressed genes from the TCGA normal and tumor datasets (Supplement file) [38]. Furthermore, using the Pearson correlation analysis, eight significant genes were investigated and visualized through the GEPIA2 module “Correlation Analysis”. To validate the analysis, the TIMER2.0 “Gene Corr” module was deployed to generate a heatmap demonstrating correlation and clinical significance. Using InteractiVenn, an online tool for analyzing interactive Venn diagrams of gene sets, a Venn diagram was illustrated to indicate the intersected genes from the binding proteins and co-expressed genes [39]. Additionally, Enrichr, a gene set enrichment analysis web-based tool, was used to perform the enrichment analysis [40]. For pathway enrichment analyses of EPS8, its binding proteins, and co-expressed genes, we employed KEGG, Reactome, and WikiPathway. For functional categories, we selected Biological Process (BP), Cellular Component (CC), and Molecular Function (MF) within the framework of Gene Ontology [41–43].

### 2.13 Statistical analysis

To understand the differential expression of EPS8 in normal and tumor tissues in TIMER2.0, the Wilcoxon test was utilized. ANOVA (Analysis of Variance) and T-tests were the primary statistical methods in UALCAN for investigating gene expression levels. Log2 transformation was used for gene expression normalization. Gene correlation analysis was done using Spearman’s correlation in GEPIA2. The statistical significance level was set at *p* < 0.05.

## 3 Results

### 3.1 Survival Analysis Data

The TCGA database was used to perform the prognostic survival analysis of EPS8 across 33 types of tumors. High expression of EPS8 is a negative prognostic marker in patients with LGG (log-rank p = 7.2e-05) and PAAD (log-rank p = 9.4e-06). However, it is a positive prognostic marker in KIRC (log-rank p = 6.1e-06) in terms of overall survival (Figure 2A, 2B, 2C). Also, in disease-free survival (DFS), high expression of EPS8 acts as a good prognostic marker in LGG (log-rank p = 0.0063) and PAAD (log-rank p = 0.0063), whereas low EPS8 expression is a good prognostic factor in KIRC (log-rank p = 7.6e-06) (Figure 2D, 2E, 2F).

**Figure 2:**
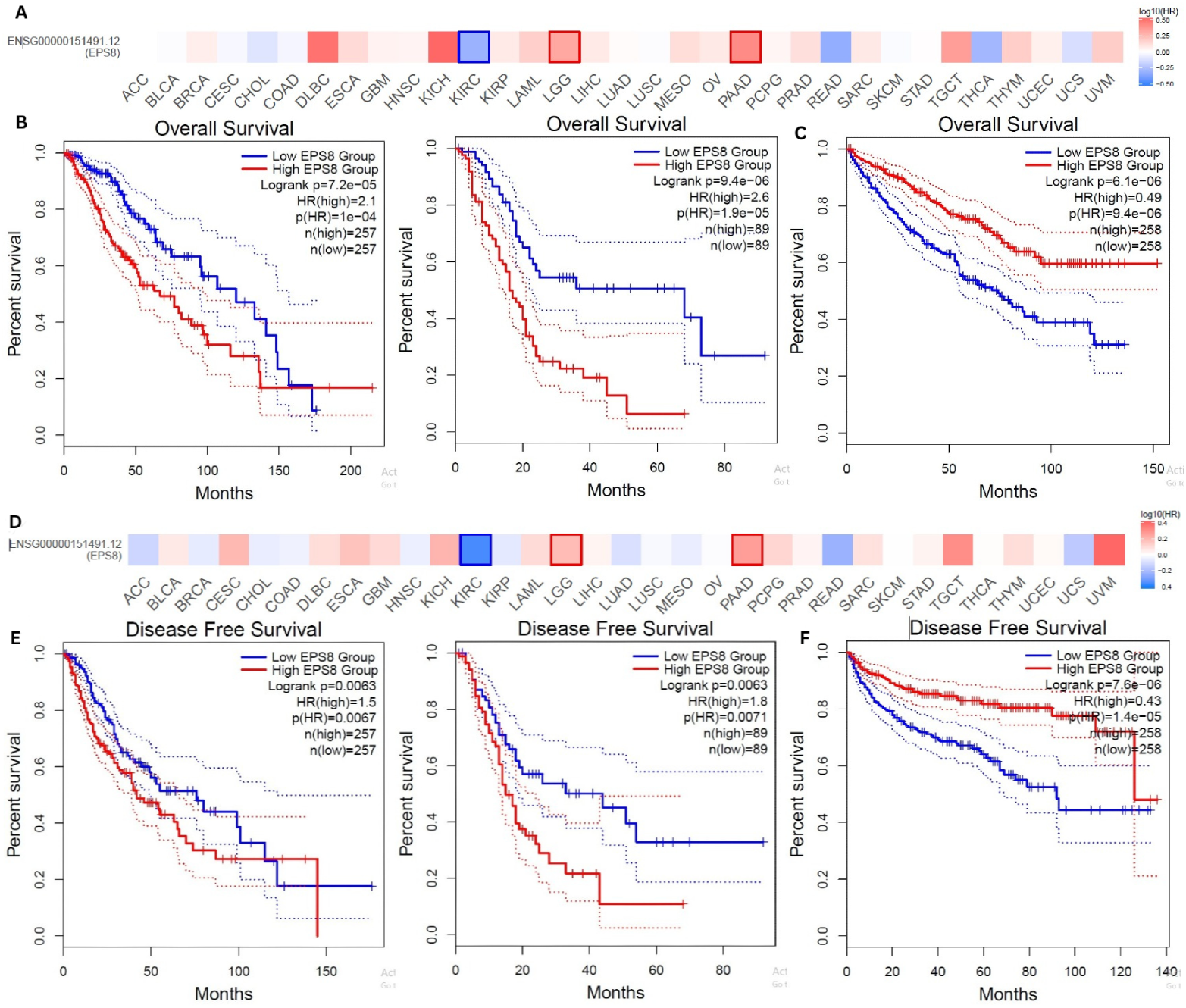
(A) The os map showing EPS8 expression in different cancers. (B)The OS plot for EPS8 expression in LGG and PAAD. (C)The OS plot for EPS8 expression in KIRC. (D) The DFS map showing EPS8 expression in different cancers. (E)The DFS plot for EPS8 expression in LGG and PAAD. (F)The DFS plot for EPS8 expression in KIRC.

### 3.2 Gene expression analysis data

To analyze the difference in EPS8 expression in tumor and normal tissues, TIMER2.0 was used. The expression level of EPS8 is noticeably higher within tumor cells than in normal tissue types, significantly in GBM, KICH, KIRC, KIRP, LIHC, LUAD, LUSC, STAD, and THCA, and it is significantly lower for BLCA, BRCA, COAD, HNSC, PRAD, READ, SKCM, and UCEC (Figure 3A). Furthermore, GEPIA2 was used to validate these results. The results of the validation indicated that EPS8 was expressed at higher levels in 20 cancer types, significantly in CHOL, ESCA, GBM, KICH, LAML, PAAD, and STAD, and was expressed at lower levels in ACC, BLCA, BRCA, CESC, OV, PRAD, TGCT, UCEC, and UCS (Supplementary File figure 1). Additionally, we applied the UALCAN database to assess the various expression levels of EPS8 across 24 types of cancer (Figure 3B). EPS8 is upregulated in 14 different types of cancer, significantly in CHOL, GBM, KICH, KIRC, KIRP, LIHC, LUAD, SKCM, STAD, THCA, and is significantly downregulated in BLCA, BRCA, COAD, PRAD, READ, and UCEC when comparing normal vs primary tumors. In particular, SKCM gene expression was significantly boosted in metastatic tumors as compared to primary tumors. Additionally, in KIRC and PAAD, EPS8 expression was found to correlate with tumor stage as determined by the GEPIA2 (*P* < 0.05)(Figure 3C).

**Figure 3:**
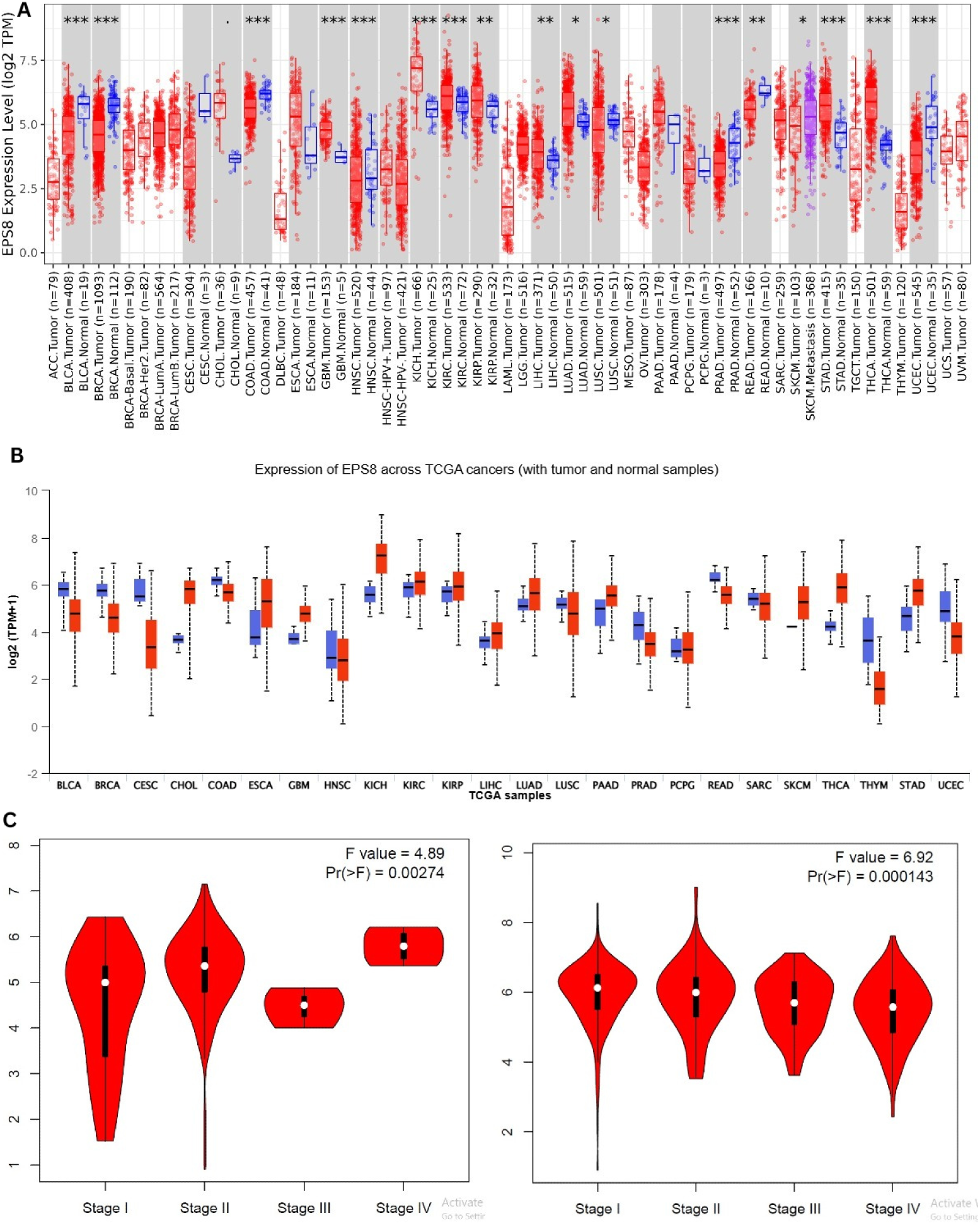
Expression analysis of EPS8 across various tumors. (A) Pan-cancer comparison of EPS8 expression between tumor and normal samples using TIMER 2.0. (B) Pan-cancer evaluation of EPS8 expression across all TCGA tumor types utilizing UALCAN. (C) Comparative analysis of EPS8 expression among different cancer stages, including KIRC and PAAD (*p* < 0.05), was conducted via GEPIA2.

### 3.3 Methylation analysis

Methylation analysis was conducted to examine the link between EPS8 DNA methylation and tumor development. EPS8 promoter methylation levels were significantly higher in KIRC and PRAD, while they were significantly lower in BLCA, BRCA, CESC, KIRP, LUAD, LUSC, TGCT, THCA, and UCEC (Supplementary File Figure 2). The visualization indicating the chromosomal distribution of methylation probes connected to EPS8 was derived from the SMART database (Figure 4A). A total of 43 probes were analyzed in order to assess the status of DNA methylation of EPS8. For the probe cg01975858 located in the Open Sea area (i.e, non-CpG island) region, we noted a prevailing trend of increased methylation in tumor tissues in comparison to normal tissue in various types of cancer, which suggests some control relevance outside promoter CpG islands. However, this pattern showed variation and was not statistically significant across multiple malignancies (Figure 4B). In the analysis of the CpG-aggregated methylation of EPS8, it was found that the EPS8 methylation level in BLCA, CHOL, COAD, ESCA, HNSC, KIRC, KIRP, LIHC, LUAD, LUSC, PAAD, READ, STAD, THCA, and UCEC is significantly low (Figure 4C).

**Figure 4:**
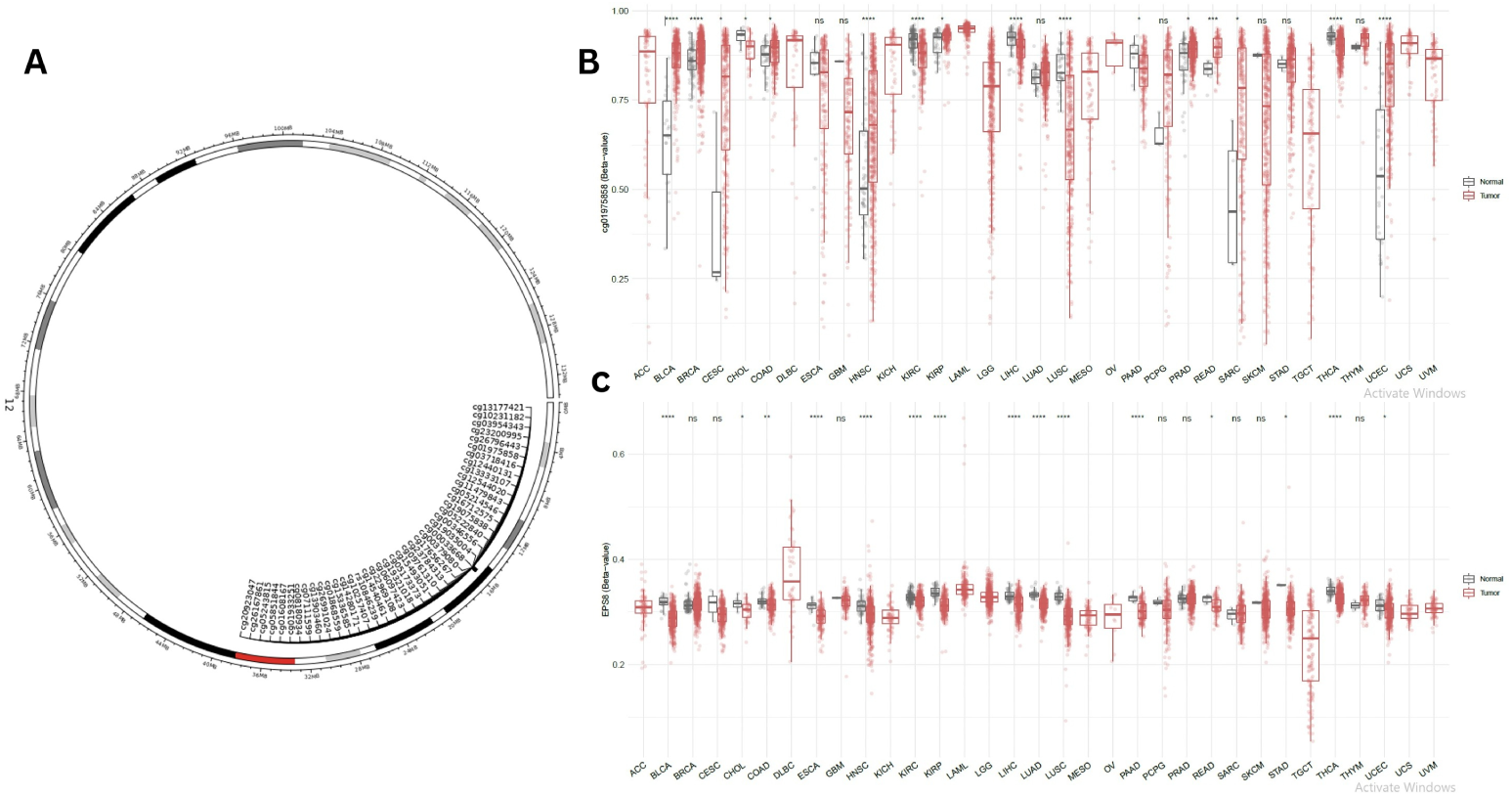
(A) Chromosomal maps of distributed methylation probes linked to EPS8. (B) Comparison of EPS8 methylation in tumor samples (red) and normal tissues (gray) among different cancer types, based on the open-sea probe cg01975858. Statistical significance was evaluated using the Wilcoxon test and denoted by asterisks: \**p* < 0.05, \*\**p* < 0.01, \*\*\**p* < 0.001, and \*\*\*\**p* < 0.0001. (C) The aggregated methylation analysis of CpG sites of EPS8 in human cancer was performed using SMART.

### 3.4 Immune infiltration assessment of EPS8

The tumor microenvironment (TME), a fundamental element in tumor prognosis, is made up of immune cells and, specifically, cancer-associated fibroblasts (CAFs), tumor-associated macrophages (TAMs), and T cells, among others [44, 45]. Figures 5A-5C show EPS8 expression and positive correlation with CAFs in BRCA, BRCA-LumA, CESC, HNSC, HNSC-HPV, LIHC, PRAD, TGCT, and THYM, and a negative correlation in ESCA, KIRP, SKCM-Metastasis, STAD, and THCA. Furthermore, T-follicular helper cells (Tfh), also a fundamental tumor-infiltrating immune cell of the TME [46, 47], demonstrated a positive correlation with EPS8 expression in UVM tumors, and KIRC tumors show a negative correlation, as noted in Figure 6: A-C.

**Figure 5:**
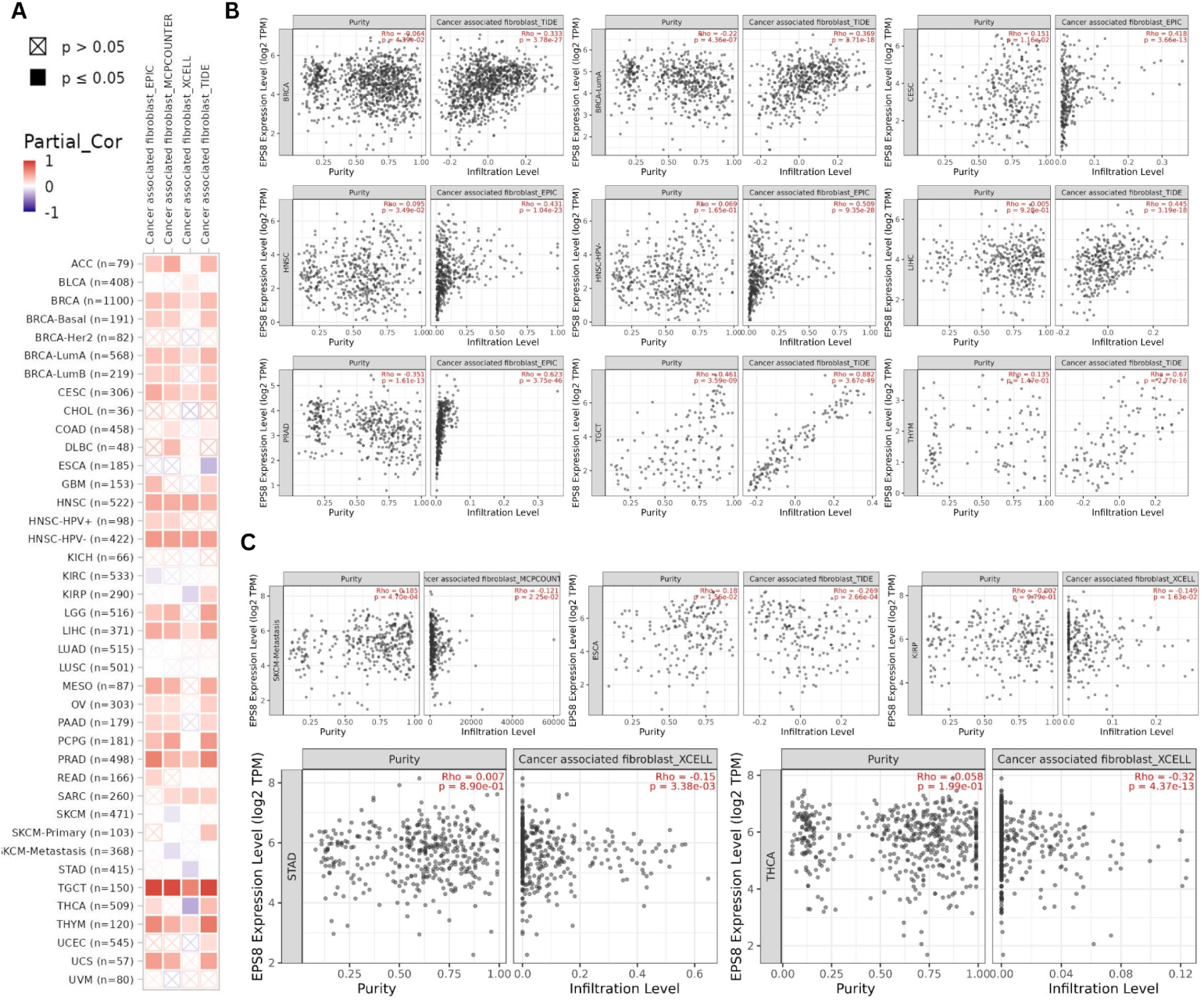
Correlation analysis of EPS8 and immune infiltration of CAF. Among the TCGA dataset, EPIC, MCPCOUNTER, XCELL, and TIDE algorithms were used to assess the correlation. A) presents the correlation heatmap between EPS8 expression and immune infiltration of CAF. B) Illustrates the scatter plot for the most positively correlated cancers. C) Showcases the scatter plot of the most negatively correlated cancers.

**Figure 6:**
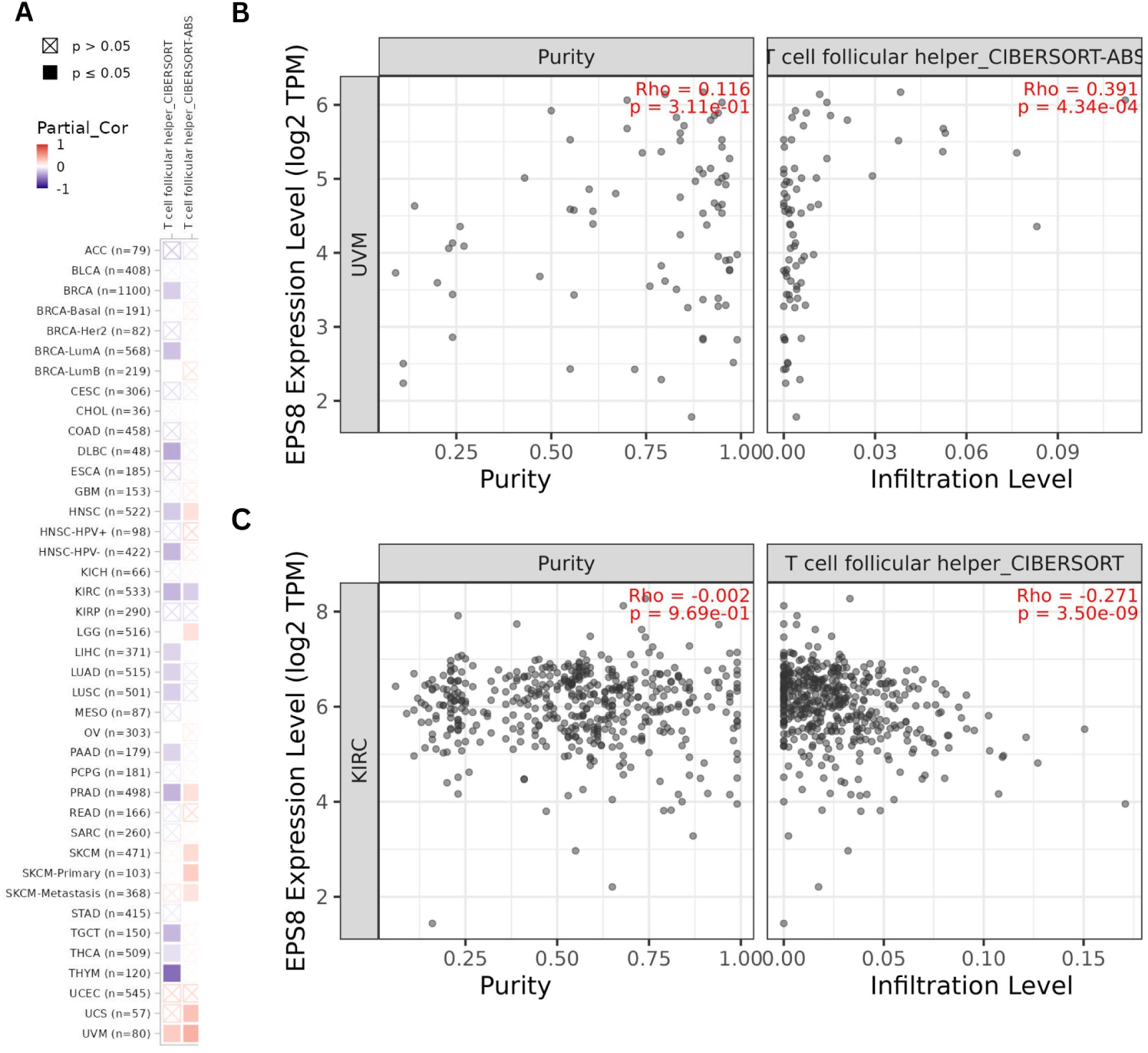
Correlation analysis of EPS8 expression and immune infiltration of Tfh cells. CIBERSORT and CIBERSORT-ABS algorithms were employed to investigate the potential correlation across the TCGA dataset. A) presents the correlation heatmap between EPS8 expression and T-follicular helper cells. B) Demonstrates the scatter plot for the most positively correlated cancers. C) Demonstrates the scatter plot for the most negatively correlated cancers.

### 3.5 Genetic alteration analysis data

The cBioportal was used to analyze genetic modifications within tumors across TCGA cohorts. EPS8 shows modification frequency peaking at approx. 6% in TGCT and UCEC. In TGCT, modifications are mostly “Amplification”, whereas UCEC predominantly shows “Mutation” alteration. In UCS, MESO, LGG, and THYM cases with genetic modifications, all show “Amplification” of EPS8. In the case of OV with alteration, “Amplification” was the most dominant, followed by “Mutation”. LUAD had a modification of every type, with “Deep deletion” being the most evident at (< 2%). Examples of “Rare Fusion” and “Multiple Alterations” are scattered across groups such as STAD and LUAD, respectively (Figure 7A). Additionally, EPS8 mutations in TCGA samples include 122 missense, 33 truncation, 5 splice, and 8 fusion mutations, with missense mutations being the most common (Figure 7B). Truncation R571* was located in the SH3 domain and was present in 1 case of LUSC and STAD and 3 cases of UCEC. EPS8 frameshift deletions have been found in BRCA, KICH, SKCM, and STAD cancers, and LUAD was identified with a frameshift insertion. The alteration P639Lfs*16 (Allan freq = 0.33, Mutation sample > 6000) was found in STAD, where Proline (P) was substituted for Leucine (L) at site 369 of the EPS8 protein. Additionally, we analyzed EPS8 genetic alterations and how they affected cancer patients’ prognosis. EPS8 mutations indicated a good prognosis for STAD and UCEC patients regarding PFS significantly. In contrast, EPS8 mutations in SKCM patients indicated a significantly worse prognosis in terms of DSS and OS (Figure 7C). Furthermore, a significantly worse prognosis in LUSC and BLCA patients in terms of DFS was also observed. However, LUSC and BLCA had relatively few samples for the altered group (Supplementary File Figure 3).

**Figure 7:**
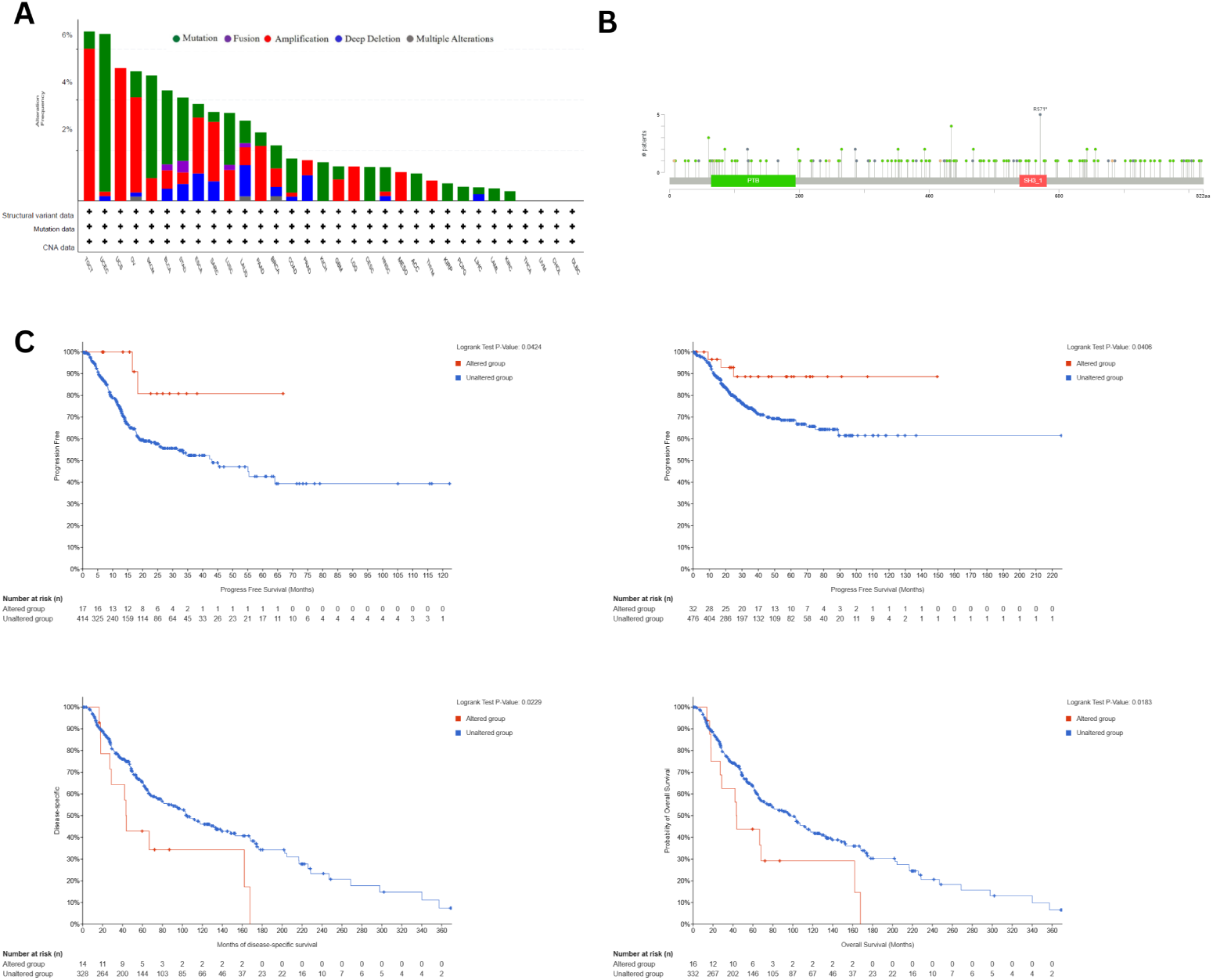
Visualization of the genetic alteration analysis of EPS8 (A) Cancer type summary for the EPS8 gene. (B) Mutation plot for EPS8. (C) Progression-free survival plot for STAD and UCEC (*p* < 0.05) (Top), Disease-specific-survival and Overall Survival curve of SKCM (Bottom) (*p* < 0.05).

### 3.6 Proteomic analysis

We utilized the UALCAN database in order to investigate the proteomic expression of EPS8. The expression of EPS8 in COAD, KIRC, KIRC (extended), LUAD, PAAD, GBM, and GBM (extended) was significantly elevated in tumor tissues. However, in BRCA, OV, UCEC, UCEC (extended), LUSC, and HNSC, significantly lower expression was observed (Figure 8B). Additionally, variations in EPS8 expression in tumor and normal tissues were observed through the statistical analysis (Supplementary File Figure 4).

**Figure 8:**
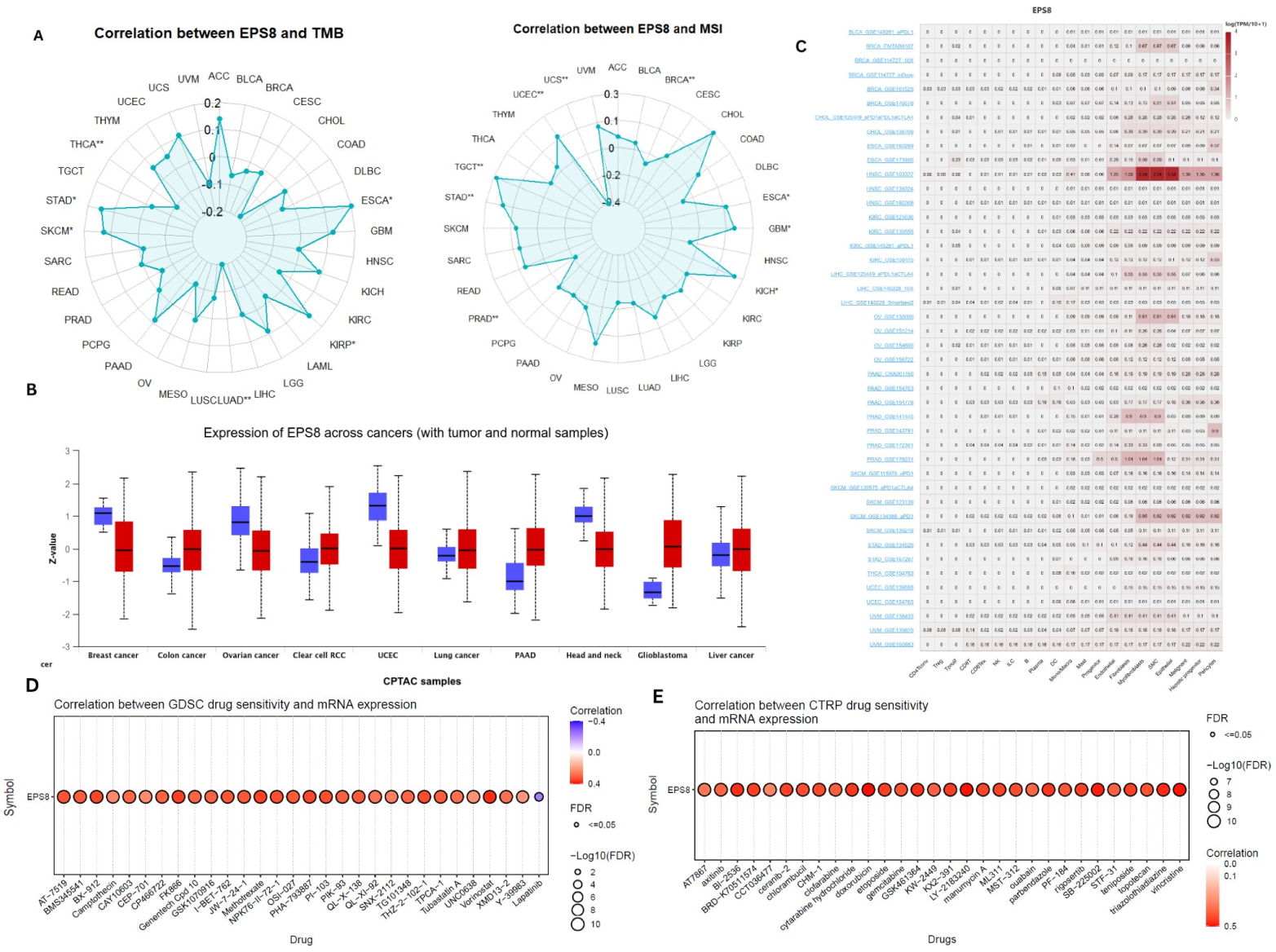
(A) EPS8 expression correlating with TMB and MSI pan-cancer context. (B) Proteomics analysis of EPS8 across various tumors utilizing UALCAN. (C) Pan-cancer Single-cell functional analysis of EPS8 from the scRNA-seq database. (D) Correlation between EPS8 mRNA expression and the top GDSC drug response. (E) Correlation between EPS8 mRNA expression and the top CRTP drug response.

### 3.7 TMB and MSI correlation analysis of EPS8

Evaluation of EPS8 expression correlation with TMB and MSI showed significant results in various cancers. EPS8 showed a significant positive connection in STAD, ESCA, SKCM, and KIRP and a negative correlation with LUAD and THCA with TMB. Additionally, EPS8 expression showed a positive correlation with MSI in STAD, TGCT, UCEC, GBM, ESCA, and KICH (\**p* < 0.05, \*\**p* < 0.01), and a significant negative correlation with MSI in PRAD, BRCA, and UCS (\**p* < 0.05, \*\**p* < 0.01) (Figure 8A). It demonstrates that EPS8 showed correlation with both TMB and MSI in STAD and ESCA significantly. Therefore, EPS8 might impact the immune response to cancer through its association with these cancers.

### 3.8 Single-cell functional analysis

We employed the TISCH2 database to understand EPS8 expression across cell lineages within the TME, which consists of immune, stromal, and cancer cells [48]. EPS8 expression was downregulated in immune cells, particularly CD4 Tconv, Treg, Tprolif, CD8T, NK, B cells, Plasma cells, ILC, and DC, compared with stromal cells like fibroblasts, epithelial cells, and myofibroblasts (Figure 8C).

### 3.9 Pan-cancer correlation with EPS8 and immune microenvironment

Immune cells play a very important role in regulating cancer development and prognosis, as well as influencing immunotherapy efficiency [49]. EPS8 was significantly negatively associated with immune scores in SKCM, BLCA, and LUSC, and it was positively related with ACC, LAML, PCPG, PRAD, LGG, and PAAD (Supplementary File Figure 5C). Furthermore, EPS8 was positively correlated with neutrophils, resting mast cells, T cells CD4 memory resting, B cells naive, and Macrophages M2, while it was negatively related to Macrophages M0, B cells memory, plasma cells, NK cells activated, T cells follicular helper, T cells CD8, and T cells regulatory Tregs in most tumor samples (Supplementary File Figure 5E). EPS8 showed positive relations with immune checkpoint-associated genes in PRAD, ACC, UVM, LGG, LAML, PAAD, THYM, KIRC, THCA, GBM, READ, OV, STAD, COAD, BRCA, and UCEC and negative associations in TGCT, BLCA, and CESC (Supplementary File Figure 5A). Almost all immune inhibitory or stimulating genes showed positive association with EPS8 in PRAD, HNSC, LIHC, LGG, UVM, OV, UCEC, KIRC, THCA, THYM, ACC, COAD, PAAD, PCPG, BRCA, STAD, GBM, LAML, READ, and SARC, and negative association in TGCT, CESC, and BLCA (Supplementary File Figure 5B, 5D).

### 3.10 Drug response analysis

Drug response analysis revealed positive correlation for several compounds from the GDSC and CTRP databases, notably, AZD8055 (mTOR Inhibitor), BMS-345541 (IKK2/IKK1 inhibitor), JQ1(BET Inhibitor), PHA-793887(CDK Inhibitor), PI-103(PI3K Inhibitor), TPCA-1(IKK-2 inhibitor), TG-101348(JAK2/FLT3 inhibitor), Vorinostat (HDAC inhibitor), Methotrexate(DHFR inhibitor), TW-37(Bcl-2 inhibitor), and OSI-027 (mTOR inhibitor). Additionally, a negative correlation was found between Trametinib (MEK inhibitor) and Selumetinib (MEK inhibitor) (Supplementary file). This reveals that EPS8 can be an important biomarker for cancer therapy.

### 3.11 EPS8 related enrichment analysis

Using the STRING and GEPIA2 databases, 50 binding proteins and 100 co-expressed genes of EPS8 were obtained. The results were combined to perform the enrichment analysis of EPS8. As shown in Figure 9A, the network represents the PPI network of EPS8. Furthermore, from the 100 most co-expressed genes, eight genes were selected for deeper examination, which showed a positive correlation with EPS8, including ARHGAP18 (*R* = 0.52), FCHO2 (*R* = 0.5), ARHGEF12 (*R* = 0.48), FAM13A (*R* = 0.47), ARHGAP42 (*R* = 0.46), ZBTB38 (*R* = 0.46), PLS1 (*R* = 0.37), and WASL (*R* = 0.41) as demonstrated in Figure 9C. The heatmap obtained from TIMER2.0 showed positive correlations among the selected eight genes and EPS8 in most tumors, as shown in Figure 9D. It can be observed from the Venn diagram intersection illustration that PLS1 and WASL are the common members of the co-expressed genes and the binding proteins, as illustrated in Figure 9B. Additionally, GO Biological Process showed the association of EPS8 with “Cellular component assembly” and “Actin filament capping”. GO cellular component analysis revealed that EPS8 was connected to “Actin-based cell projection” and “Actin cytoskeleton”. Furthermore, GO molecular functions demonstrated a significant linkage with “Cadherin binding” and “Actin binding”. It is observed by the Reactome 2024 pathway that EPS8 contributes to “Sensory Processing of Sound” with *p*-value < 1 × 10^−33^ and odds ratio > 80. The WikiPathway 2024 Human revealed that EPS8 is connected with “Regulation of Actin Cytoskeleton” and “EGF-EGFR Signaling” with *p*-value < 1 × 10^−6^ and odds ratio > 10. The KEGG 2026 enrichment analysis disclosed that EPS8 was connected with “Regulation of Actin Cytoskeleton”, and “Adherens Junction” with *p*-value < 1 × 10^−7^ and odds ratio > 10, as shown in Figure 10A-F. The results showing the most significance according to p-values: Reactome Pathways and GO Cellular Component are demonstrated in table 1 and 2, respectively. The rest if the tables are provided in the supplementary file Table 1-4.

**Figure 9:**
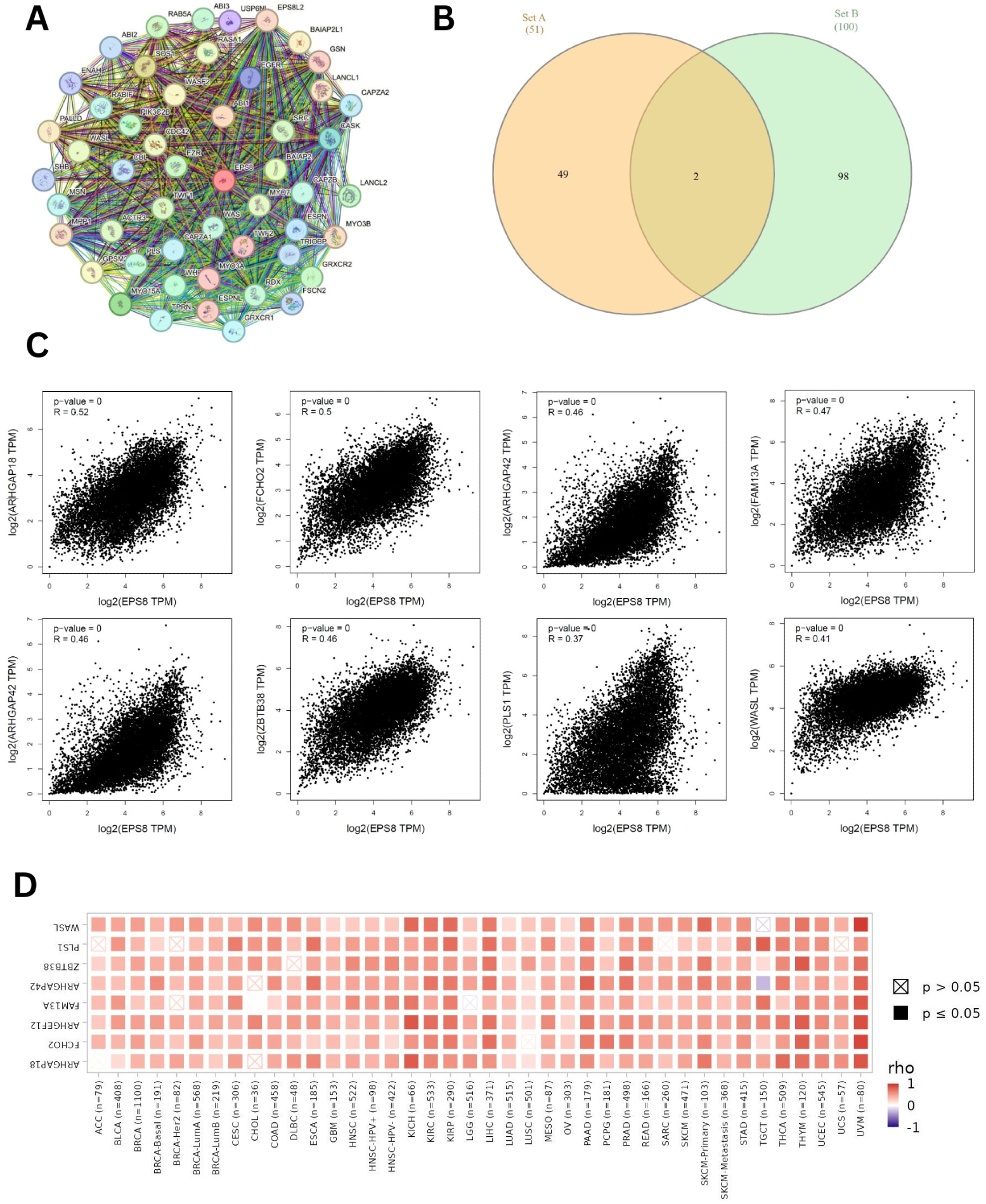
EPS8-related genes analysis visualization. (A) PPI network using STRING. (B) An Intersection of the EPS8 binding proteins and correlated genes, illustrated using a Venn diagram. (C) The scatter plots demonstrating the correlation between EPS8 expression and eight co-expressed genes (ARHGAP18, FCHO2, ARHGEF12, FAM13A, ARHGAP42, ZBTB38, PLS1, and WASL). (D) Heatmap illustrating the linkage between EPS8 expression and eight selected genes in TCGA cancer.

**Figure 10:**
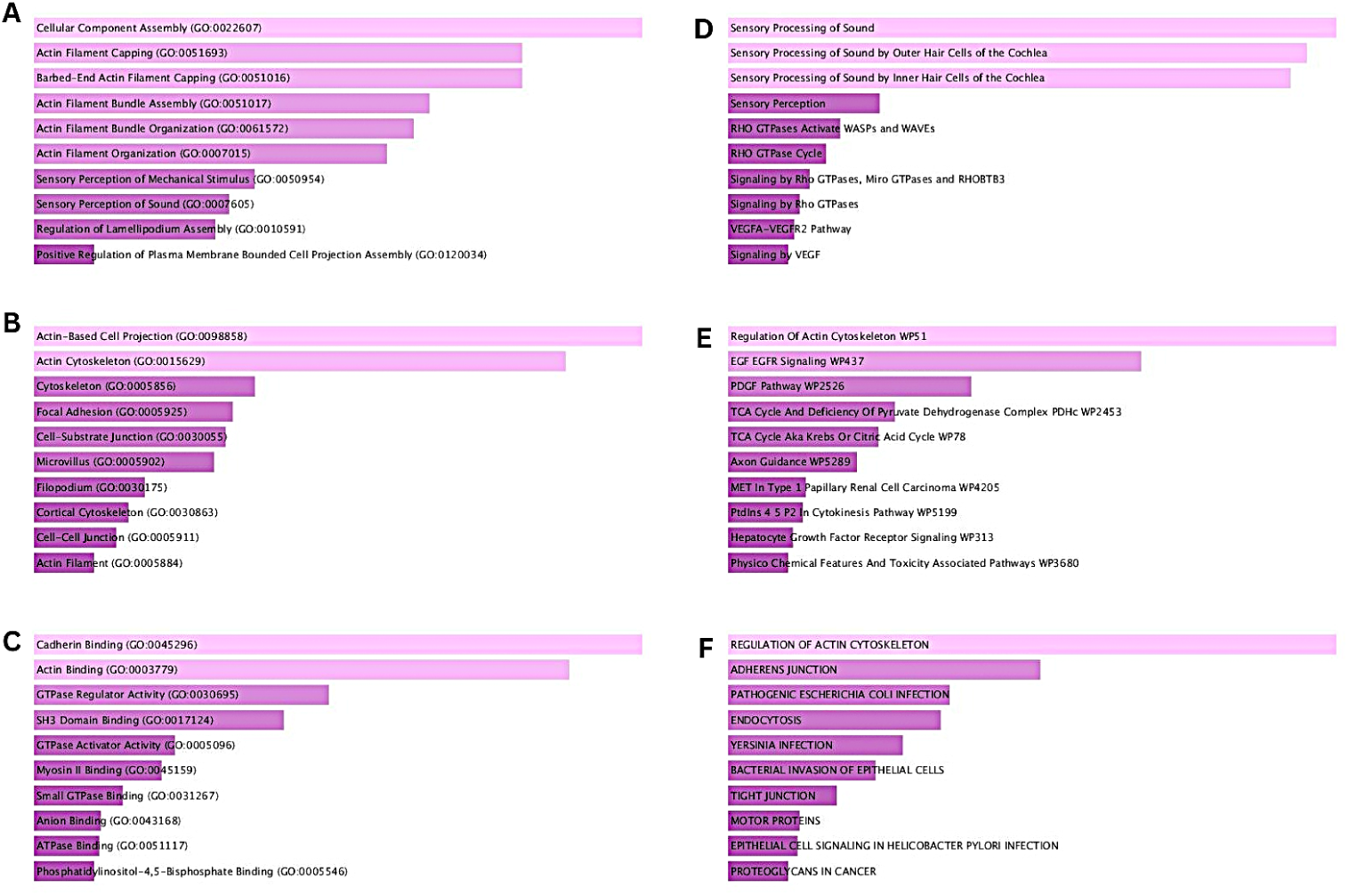
Gene enrichment analysis of EPS8 using Enricher. The bar charts indicate the relationship between EPS8 and multiple pathways, and higher significance is represented by a lighter color. (A) GO Biological Process 2025, (B) GO Cellular Component 2025, (C) GO Molecular Function 2025, (D) Reactome Pathways 2024, (E) WikiPathway 2024 Human, and (F) KEGG 2026.

**Table 1:**
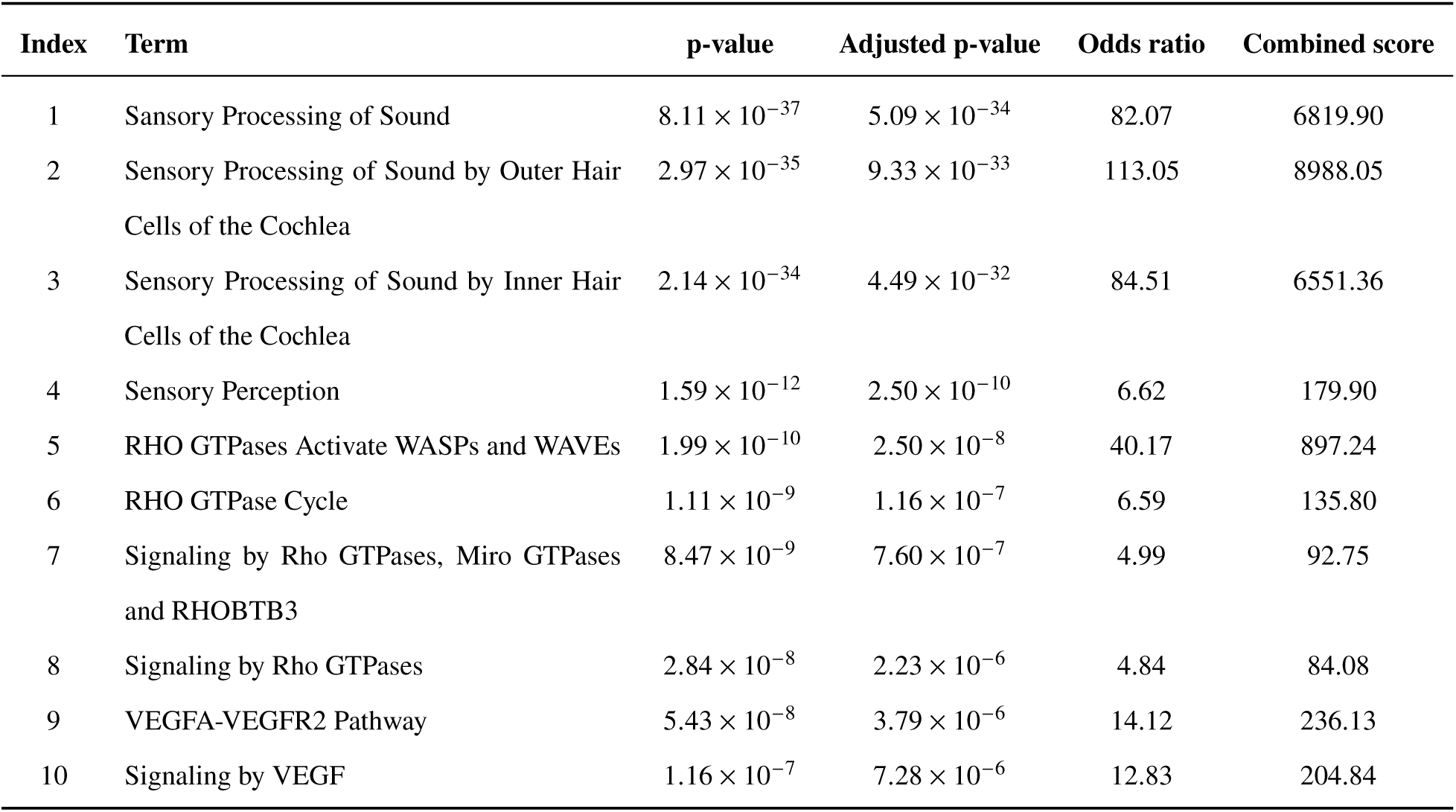
Reactome Pathways 2024 enrichment analysis.

| Index | Term | p-value | Adjusted p-value | Odds ratio | Combined score |
| --- | --- | --- | --- | --- | --- |
| 1 | Sansory Processing of Sound | $8.11 \times 10^{-37}$ | $5.09 \times 10^{-34}$ | 82.07 | 6819.90 |
| 2 | Sensory Processing of Sound by Outer Hair Cells of the Cochlea | $2.97 \times 10^{-35}$ | $9.33 \times 10^{-33}$ | 113.05 | 8988.05 |
| 3 | Sensory Processing of Sound by Inner Hair Cells of the Cochlea | $2.14 \times 10^{-34}$ | $4.49 \times 10^{-32}$ | 84.51 | 6551.36 |
| 4 | Sensory Perception | $1.59 \times 10^{-12}$ | $2.50 \times 10^{-10}$ | 6.62 | 179.90 |
| 5 | RHO GTPases Activate WASPs and WAVES | $1.99 \times 10^{-10}$ | $2.50 \times 10^{-8}$ | 40.17 | 897.24 |
| 6 | RHO GTPase Cycle | $1.11 \times 10^{-9}$ | $1.16 \times 10^{-7}$ | 6.59 | 135.80 |
| 7 | Signaling by Rho GTPases, Miro GTPases and RHOBTB3 | $8.47 \times 10^{-9}$ | $7.60 \times 10^{-7}$ | 4.99 | 92.75 |
| 8 | Signaling by Rho GTPases | $2.84 \times 10^{-8}$ | $2.23 \times 10^{-6}$ | 4.84 | 84.08 |
| 9 | VEGFA-VEGFR2 Pathway | $5.43 \times 10^{-8}$ | $3.79 \times 10^{-6}$ | 14.12 | 236.13 |
| 10 | Signaling by VEGF | $1.16 \times 10^{-7}$ | $7.28 \times 10^{-6}$ | 12.83 | 204.84 |

**Table 2:** GO Cellular Component 2025 enrichment analysis.

| Index | Term | p-value | Adjusted p-value | Odds ratio | Combined score |
| --- | --- | --- | --- | --- | --- |
| 1 | Actin-Based Cell Projection (GO:0098858) | $1.173 \times 10^{-16}$ | $1.724 \times 10^{-14}$ | 30.33 | 1112.49 |
| 2 | Actin Cytoskeleton (GO:0015629) | $3.097 \times 10^{-15}$ | $2.276 \times 10^{-13}$ | 10.54 | 352.06 |
| 3 | Cytoskeleton (GO:0005856) | $1.895 \times 10^{-9}$ | $9.286 \times 10^{-8}$ | 5.22 | 104.79 |
| 4 | Focal Adhesion (GO:0005925) | $4.924 \times 10^{-9}$ | $1.810 \times 10^{-7}$ | 6.78 | 129.71 |
| 5 | Cell-Substrate Junction (GO:0030055) | $6.677 \times 10^{-9}$ | $1.963 \times 10^{-7}$ | 6.63 | 124.89 |
| 6 | Microvillus (GO:0005902) | $1.099 \times 10^{-8}$ | $2.692 \times 10^{-7}$ | 22.47 | 411.78 |
| 7 | Filopodium (GO:0030175) | $2.140 \times 10^{-7}$ | $4.494 \times 10^{-6}$ | 19.52 | 299.81 |
| 8 | Cortical Cytoskeleton (GO:0030863) | $4.315 \times 10^{-7}$ | $7.928 \times 10^{-6}$ | 17.43 | 255.38 |
| 9 | Cell-Cell Junction (GO:0005911) | $7.299 \times 10^{-7}$ | $1.192 \times 10^{-5}$ | 6.17 | 87.14 |
| 10 | Actin Filament (GO:0005884) | $1.877 \times 10^{-6}$ | $2.759 \times 10^{-5}$ | 13.73 | 181.09 |

## 4 Discussion

The TCGA Pan-Cancer dataset comprises approximately 11,000 tumors across 33 cancer types with multiomics data, providing a valuable resource for investigating the oncogenic mechanisms underlying cancer-associated genes [50]. EPS8 acts as a signaling adaptor that regulates multiple cellular protrusions through actin cytoskeleton dynamics. Together with SOS1 and ABI1, it forms a trimeric complex that facilitates the conversion of Ras to Rac [51]. According to the Ensembl database, EPS8 is an epidermal growth factor receptor (EGFR) substrate located on chromosome 12 (15,620,134–15,882,330) on the reverse strand of the GRCh38 assembly. It contains 50 transcripts, 218 orthologs, 3 paralogs, and is associated with 2 phenotypes, geneID: ENSG00000151491.15 [52].

Although previous studies have suggested a significant role for EPS8 in tumor development [53], its significance has not been comprehensively explored in the pan-cancer context. Therefore, we performed a comprehensive pan-cancer analysis to characterize the molecular, prognostic, and immunological significance of EPS8 across multiple cancer types.

Our results showed that elevated EPS8 expression was associated with poor overall survival (OS) and disease-free survival (DFS) in patients with LGG and PAAD, whereas it was associated with a favorable prognosis in patients with KIRC. Additionally, EPS8 expression was significantly higher in metastatic SKCM than in primary tumors, suggesting a potential association with tumor progression and metastasis. Collectively, these findings support the prognostic relevance of EPS8 across multiple cancer types and warrant further investigation into its biological role in cancer progression.

DNA methylation involves the addition of a methyl group to cytosines at CpG sites. Cancer is characterized by a pattern of global hypomethylation at CpG islands, making DNA methylation a valuable biomarker for early detection and prognosis [54]. Our study revealed that the promoter methylation level of EPS8 was significantly decreased across most cancer types. Additionally, the probe cg01975858, located in the Open Sea region, exhibited increased methylation in most malignancies examined, suggesting a potential regulatory role outside promoter CpG islands. Collectively, these findings indicate that EPS8 may be involved in epigenetic regulation across multiple cancer types.

Immune infiltration analysis provides insights into tumor prognosis, identifies immune-hot tumors, and links immune cell composition with mutation burden [55]. Cancer-associated fibroblasts (CAFs) have been associated with poor prognosis, chemotherapy resistance, and disease recurrence [30]. In our study, EPS8 expression was positively correlated with CAF infiltration in multiple cancers, including BRCA and BRCA-LumA, and negatively correlated with SKCM-Metastasis, among others. Additionally, T follicular helper (Tfh) cells promote germinal center formation, sustain CD8^+^ T-cell proliferation, and recruit CD8^+^ T cells into tumors [56]. We observed a positive correlation between EPS8 expression and Tfh cell infiltration in UVM and a negative correlation in KIRC. These findings suggest that EPS8 may contribute to the regulation of tumor-infiltrating immune cells and, consequently, influence tumor prognosis. Overall, the observed associations indicate that EPS8 is closely linked to immune cell infiltration and may play a role in regulating the tumor immune microenvironment, supporting its potential as a biomarker for future immunotherapy-related research.

Cancer is driven by the accumulation of genetic and epigenetic alterations in oncogenes [57]. Comprehensive profiling of these alterations provides valuable diagnostic, prognostic, and therapeutic information across multiple tumor types [58]. Our genetic alteration analysis revealed that TGCT and UCEC exhibited the highest alteration frequencies for EPS8, with amplification being the most prevalent alteration. Furthermore, analysis of the clinical impact of EPS8 alterations showed that patients with SKCM harboring EPS8 alterations had poorer overall survival (OS) and disease-free survival (DFS). These findings suggest that EPS8 alterations are associated with cancer prognosis and highlight their potential clinical relevance.

Immunotherapy has emerged as an effective treatment strategy for many cancers, with tumor mutational burden (TMB) and microsatellite instability (MSI) serving as established biomarkers for predicting immune checkpoint inhibitor (ICI) response [59]. Correlation analysis revealed that EPS8 expression was significantly associated with TMB in six cancer types and with MSI in nine cancer types. Notably, STAD and ESCA showed significant correlations with both TMB and MSI. These findings suggest that EPS8 may be associated with immunotherapy-related biomarkers in these cancers and warrant further investigation into its potential clinical relevance.

Single-cell analysis measures gene expression at the individual cell level, enabling the identification of cancer subclones and diverse cellular states [60]. Our analysis revealed that EPS8 was more highly expressed in stromal cells than in immune cells. Previous studies have suggested that lower gene expression in immune cells may reflect impaired anti-tumor immunity [61,62]. Accordingly, EPS8 may influence immune responses within the tumor microenvironment (TME) and represents a potential biomarker for further investigation.

The TME consists of immune cells, stromal cells, and endothelial cells that collectively influence tumor progression, invasion, and metastasis [63]. Our analysis showed that EPS8 expression was significantly associated with the immune score in three cancer types and positively correlated with the stromal score in six cancer types. Additionally, EPS8 expression was positively correlated with M2 macrophages and neutrophils, and negatively correlated with CD8^+^ T cells, activated NK cells, and regulatory T cells (Tregs), among others. Neutrophil infiltration has been associated with shorter DFS, DSS, and OS, as well as an increased risk of mortality [64]. Similarly, M2 macrophages promote angiogenesis, lymphangiogenesis, and immunosuppression [65, 66], whereas high CD8^+^ T-cell infiltration is generally associated with improved OS and DFS [67]. Regulatory T cells have also been implicated in suppressing anti-tumor immunity and influencing responses to immunotherapy [33]. Furthermore, immune checkpoint inhibitors (ICIs) have become a cornerstone of modern cancer immunotherapy by blocking inhibitory receptors that suppress T-cell activity [68]. Clinical studies have demonstrated the efficacy of ICIs in several cancers, including melanoma and NSCLC [68, 69]. Our analysis further showed that EPS8 expression was positively correlated with immune checkpoint genes in 16 cancer types and negatively correlated in 3 cancer types. Collectively, these findings suggest that EPS8 is associated with immune regulation across multiple cancers and support further investigation of its potential role in the tumor immune microenvironment and immunotherapy-related research.

Drug sensitivity analysis provides insights into the relationship between gene expression and drug resistance, sensitivity, and dependency [70]. Our pan-cancer analysis revealed that EPS8 expression was positively correlated with sensitivity to AZD8055 and OSI-027. Both AZD8055 and OSI-027 inhibit mTOR signaling, a central pathway regulating cell growth and survival, thereby suppressing cell proliferation, disrupting the cell cycle, and promoting tumor cell death [71]. Additionally, EPS8 expression was negatively correlated with Trametinib and Selumetinib. Trametinib inhibits the MAPK/ERK signaling pathway, reducing cell proliferation, disrupting the cell cycle, and promoting apoptosis [72]. These findings suggest that EPS8 expression is associated with the sensitivity of several targeted therapies and may have potential as a biomarker for guiding personalized treatment strategies.

Gene enrichment analysis in pan-cancer reveals gene-associated cancer hallmarks and provides insights into tumorigenesis, prognosis, and genome instability [73]. To identify key genes associated with EPS8, binding proteins and co-expressed genes were integrated. We found that ARHGAP18, FCHO2, ARHGEF12, FAM13A, ARHGAP42, and ZBTB38 were positively correlated with EPS8, while PLS1 and WASL were present in both gene sets. GO Biological Process analysis showed that EPS8 was significantly associated with actin filament organization and assembly, as well as cellular component assembly. These processes regulate cell shape, motility, proliferation, and invasion [74]. GO Cellular Component analysis indicated that EPS8 was primarily localized to actin cytoskeleton structures. Previous studies have shown that EPS8 plays an important role in pancreatic cancer by regulating actin cytoskeleton dynamics that influence cell shape, migration, and invasion [53]. In the GO Molecular Function category, EPS8 showed significant associations with actin binding, protein interaction functions, and Rho GTPase signaling. Actin and actin-binding proteins (ABPs) are known to regulate tumor migration, cell proliferation, invasion, and metastasis [74]. Reactome 2024 enrichment analysis further showed that EPS8 was strongly associated with sensory processing pathways, with a combined score approaching 9000. Previous studies suggest that these pathways contribute to cancer progression and immune responses [75]. Enrichment of sensory processing pathways related to cochlear hair cells was also observed, consistent with previous findings that EPS8 regulates actin bundle organization in specialized sensory cells [76]. In contrast, pathway analyses using WikiPathways 2024 Human and KEGG 2026 primarily highlighted EPS8 involvement in cytoskeleton regulation and cancer-related signaling pathways, including regulation of the actin cytoskeleton, PDGF signaling, and EGF-EGFR signaling. These pathways are closely interconnected in cancer progression, influencing cell adhesion, migration, proliferation, and survival [**?**]. Collectively, these findings suggest that EPS8 may contribute to cancer progression through the regulation of cytoskeletal dynamics, cell adhesion, and cancer-related signaling pathways involved in tumor cell migration, invasion, and proliferation, warranting further investigation as a potential therapeutic target.

Despite these findings, this study has several limitations. The analyses were primarily based on TCGA datasets, and variability in data collection, processing, and annotation across databases may introduce systematic bias. In addition, the limited sample size for certain rare tumor types may have affected the robustness of some analyses. Therefore, the findings may not be fully generalizable to all populations. Furthermore, this study relied exclusively on bioinformatics analyses without experimental validation. Future in vitro and in vivo studies are required to validate these findings and further elucidate the molecular and cellular roles of EPS8 in carcinogenesis.

## 5 Conclusion

In this study, through the pan-cancer analysis of EPS8, a wide range of characteristics of EPS8 in various cancer types are demonstrated. It is suggested by survival prognosis suggests that EPS8 may serve as a potential biomarker for LGG and PAAD. Methylation analysis suggests some control relevance outside promoter CpG islands. EPS8 manifested a notable correlation with TMB and MSI in ESCA and STAD. Higher EPS8 expression is revealed in stromal cells than in immune cells in various types of tumors by single-cell analysis. The results of the drug sensitivity analysis pointed out that EPS8 expression may act as a key factor in the prospective advancement of selected immunotherapies. EPS8-related gene enrichment analysis provides potential mechanisms through which the cell cycle pathway and diverse cellular functions in cancer are regulated by EPS8. Overall, this study provides the first comprehensive pan-cancer analysis of EPS8 by integrating multi-omics and immune-related analyses, highlighting its oncogenic, prognostic, and immunological significance across human cancers. Combined together, these results strengthen the position of EPS8 as more than a cancer-specific marker. It also points toward actin cytoskeleton regulation as a mechanistic axis worth targeting therapeutically. Further experimental and clinical research is needed to assess the practical implementation of EPS8 in cancer therapy and prognosis prediction. Future studies should validate these findings through functional in vitro and in vivo assays, as well as independent clinical cohort validation, to confirm the biological and clinical utility of EPS8 as a biomarker and therapeutic target.

## Supporting information

EPS8_Similar_Genes_Supplement

EPS8_CTRP_Significant_Corr_Supplement

EPS8_GDSC_Significant_Corr_Supplement

## CRediT authorship contribution statement

Adiba Juoairia: Concept, Data curation, Methodology, Software, Formal analysis, Result interpretation, Visualization, Validation, Supervision, Writing, Review and editing. Ayesha Akter: Formal analysis, Result interpretation, Visualization, Validation, Supervision, Writing, Review and editing. Rawaz Jahan Nima: Supervision, Writing, Review and editing. Md Azizul Haque: Supervision, Writing, Review and editing.

## Data availability

All the data used and generated are available at: GEPIA2, SMART, TIMER2.0, UALCAN, cBioPortal, Enrichr, Ensembl, GSCA, TISCH2, STRING, InteractiVenn, and the TCGAplot (v8.0.0) R package analysis (TCGAplot). All data and images generated and used in this study are included in the article and the supplementary files.

## Conflict of interest

The authors declare no competing interests.

## Funding

This research received no external funding.

## Ethics approval

The study used publicly available datasets, and all permissions were obtained by the original publishers following international guidelines.

## Acknowledgments

The authors would like to thank Ismail Hossen for their support.

## Additional Information

Supplementary Information is included with the manuscript.

## Notes

### Competing Interest Statement

The authors have declared no competing interest.

