## Supplement file for "Comprehensive Pan-Cancer Analysis Reveals the Prognostic, Molecular, and Immunological Significance of EPS8"

Table 1:GO_Biological_Process_2025_table

| **Index** | **Term** | **p-value** | **Adjusted p-value** | **Odds ratio** | **Combined score** |
| --- | --- | --- | --- | --- | --- |
| 1 | Cellular Component Assembly (GO:0022607) | 2.800e-12 | 3.201e-9 | 10.39 | 276.49 |
| 2 | Actin Filament Capping (GO:0051693) | 2.019e-11 | 7.692e-9 | 97.81 | 2408.60 |
| 3 | Barbed-End Actin Filament Capping (GO:0051016) | 2.019e-11 | 7.692e-9 | 97.81 | 2408.60 |
| 4 | Actin Filament Bundle Assembly (GO:0051017) | 9.311e-11 | 2.661e-8 | 45.00 | 1039.26 |
| 5 | Actin Filament Bundle Organization (GO:0061572) | 1.210e-10 | 2.766e-8 | 43.26 | 987.91 |
| 6 | Actin Filament Organization (GO:0007015) | 1.880e-10 | 3.581e-8 | 12.99 | 290.92 |
| 7 | Sensory Perception of Mechanical Stimulus (GO:0050954) | 1.662e-9 | 2.714e-7 | 17.34 | 350.62 |
| 8 | Sensory Perception of Sound (GO:0007605) | 2.535e-9 | 3.622e-7 | 16.53 | 327.26 |
| 9 | Regulation of Lamellipodium Assembly (GO:0010591) | 3.183e-9 | 4.042e-7 | 39.09 | 764.88 |
| 10 | Positive Regulation of Plasma Membrane-Bounded Cell Projection Assembly (GO:0120034) | 2.343e-8 | 2.491e-6 | 15.69 | 275.67 |

Table 2: GO_Molecular_Function_2025_table

| **Index** | **Term** | **p-value** | **Adjusted p-value** | **Odds ratio** | **Combined score** |
| --- | --- | --- | --- | --- | --- |
| 1 | Cadherin Binding (GO:0045296) | 2.38e-10 | 5.09e-08 | 8.39 | 185.99 |
| 2 | Actin Binding (GO:0003779) | 1.52e-09 | 1.62e-07 | 10.81 | 219.52 |
| 3 | GTPase Regulator Activity (GO:0030695) | 6.74e-07 | 4.81e-05 | 5.32 | 75.61 |
| 4 | SH3 Domain Binding (GO:0017124) | 2.11e-06 | 1.13e-04 | 18.47 | 241.33 |
| 5 | GTPase Activator Activity (GO:0005096) | 3.32e-05 | 1.42e-03 | 6.16 | 63.53 |
| 6 | Myosin II Binding (GO:0045159) | 4.68e-05 | 1.67e-03 | 58.25 | 580.74 |
| 7 | Small GTPase Binding (GO:0031267) | 1.25e-04 | 3.84e-03 | 5.84 | 52.46 |
| 8 | Anion Binding (GO:0043168) | 2.18e-04 | 5.44e-03 | 3.97 | 33.44 |
| 9 | ATPase Binding (GO:0051117) | 2.29e-04 | 5.44e-03 | 9.95 | 83.46 |
| 10 | Phosphatidylinositol-4,5-Bisphosphate Binding (GO:0005546) | 2.59e-04 | 5.54e-03 | 9.67 | 79.89 |

Table 3: KEGG_2026_table

| **Index** | **Term** | **p-value** | **Adjusted p-value** | **Odds ratio** | **Combined score** |
| --- | --- | --- | --- | --- | --- |
| 1 | REGULATION OF ACTIN CYTOSKELETON | 1.54e-11 | 2.78e-09 | 11.14 | 277.47 |
| 2 | ADHERENS JUNCTION | 2.85e-08 | 2.58e-06 | 15.31 | 266.02 |
| 3 | PATHOGENIC ESCHERICHIA COLI INFECTION | 2.85e-07 | 1.62e-05 | 8.38 | 126.33 |
| 4 | ENDOCYTOSIS | 3.58e-07 | 1.62e-05 | 7.28 | 108.05 |
| 5 | YERSINIA INFECTION | 9.39e-07 | 3.40e-05 | 9.83 | 136.40 |
| 6 | BACTERIAL INVASION OF EPITHELIAL CELLS | 1.88e-06 | 5.66e-05 | 13.73 | 181.09 |
| 7 | TIGHT JUNCTION | 5.03e-06 | 1.30e-04 | 7.91 | 96.52 |
| 8 | MOTOR PROTEINS | 1.30e-05 | 2.75e-04 | 6.99 | 78.62 |
| 9 | EPITHELIAL CELL SIGNALING IN HELICOBACTER PYLORI INFECTION | 1.37e-05 | 2.75e-04 | 12.97 | 145.28 |
| 10 | PROTEOGLYCANS IN CANCER | 1.73e-05 | 3.13e-04 | 6.72 | 73.72 |

Table 4: WikiPathways_2024_Human_table

| **Index** | **Term** | **p-value** | **Adjusted p-value** | **Odds ratio** | **Combined score** |
| --- | --- | --- | --- | --- | --- |
| 1 | Regulation Of Actin Cytoskeleton WP51 | 6.57e-12 | 1.81e-09 | 15.03 | 387.08 |
| 2 | EGF EGFR Signaling WP437 | 2.61e-09 | 3.60e-07 | 11.74 | 232.06 |
| 3 | PDGF Pathway WP2526 | 4.81e-07 | 4.43e-05 | 24.46 | 355.75 |
| 4 | TCA Cycle And Deficiency Of Pyruvate Dehydrogenase Complex PDHc WP2453 | 5.02e-06 | 3.47e-04 | 45.61 | 556.48 |
| 5 | TCA Cycle Aka Krebs Or Citric Acid Cycle WP78 | 8.35e-06 | 4.61e-04 | 39.09 | 457.08 |
| 6 | Axon Guidance WP5289 | 1.61e-05 | 7.41e-04 | 12.58 | 138.81 |
| 7 | MET In Type 1 Papillary Renal Cell Carcinoma WP4205 | 7.77e-05 | 2.93e-03 | 12.73 | 120.46 |
| 8 | PtdIns 4 5 P2 In Cytokinesis Pathway WP5199 | 8.49e-05 | 2.93e-03 | 45.30 | 424.68 |
| 9 | Hepatocyte Growth Factor Receptor Signaling WP313 | 1.15e-04 | 3.34e-03 | 18.23 | 165.27 |
| 10 | Physicochemical Features And Toxicity Associated Pathways WP3680 | 1.33e-04 | 3.34e-03 | 11.26 | 100.54 |

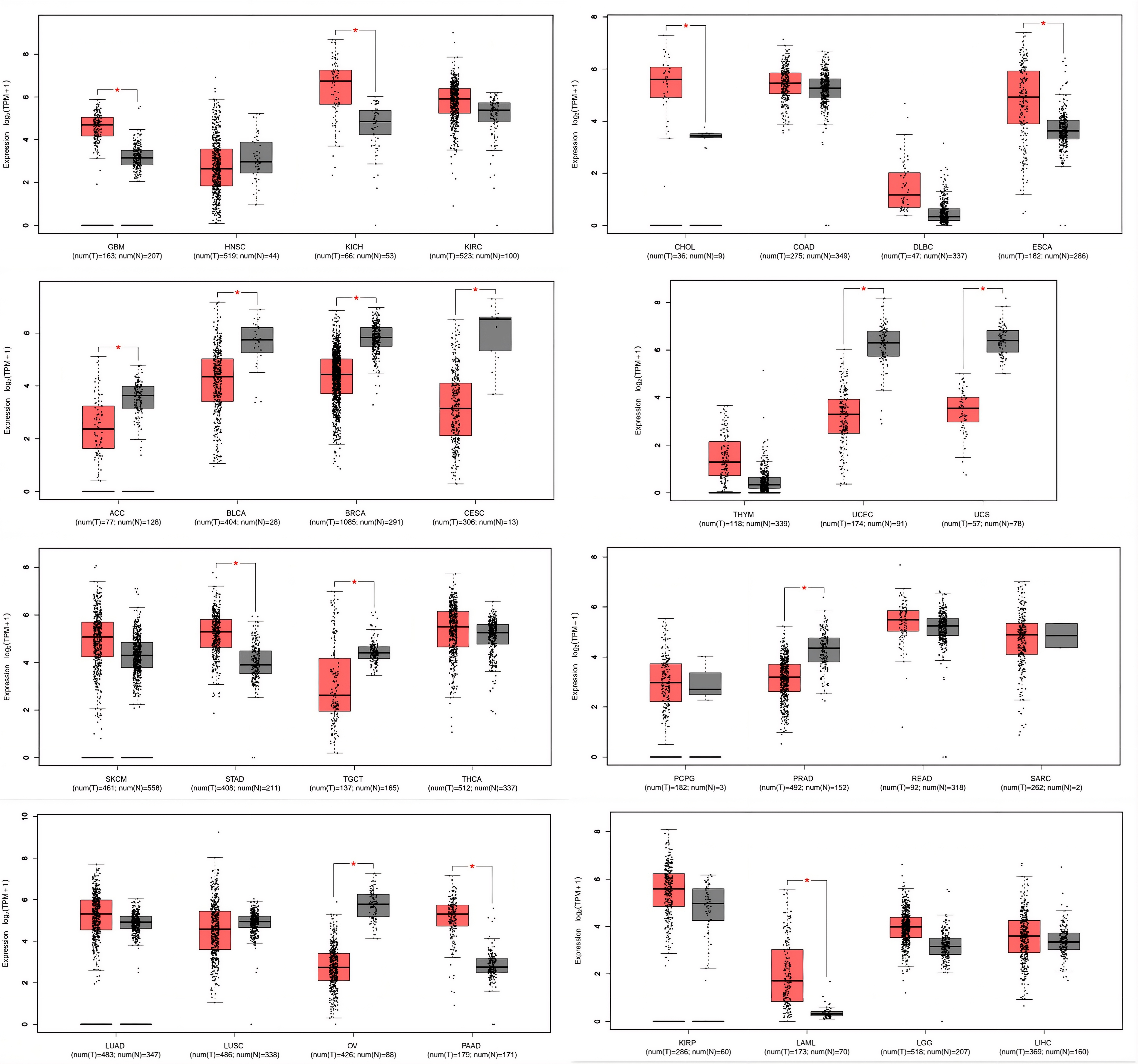

**Figure 1:** Gepia 2 was used to conduct and validate the expression analysis. Red indicates tumor tissue expression, while green indicates normal tissue expression. We can see that EPS8 was overexpressed in CHOL, ESCA, GBM, KICH, LAML, PAAD, STAD, and was expressed underexpressed in ACC, BLCA, BRCA, CESC, OV, PRAD, TGCT, UCEC, and UCS.

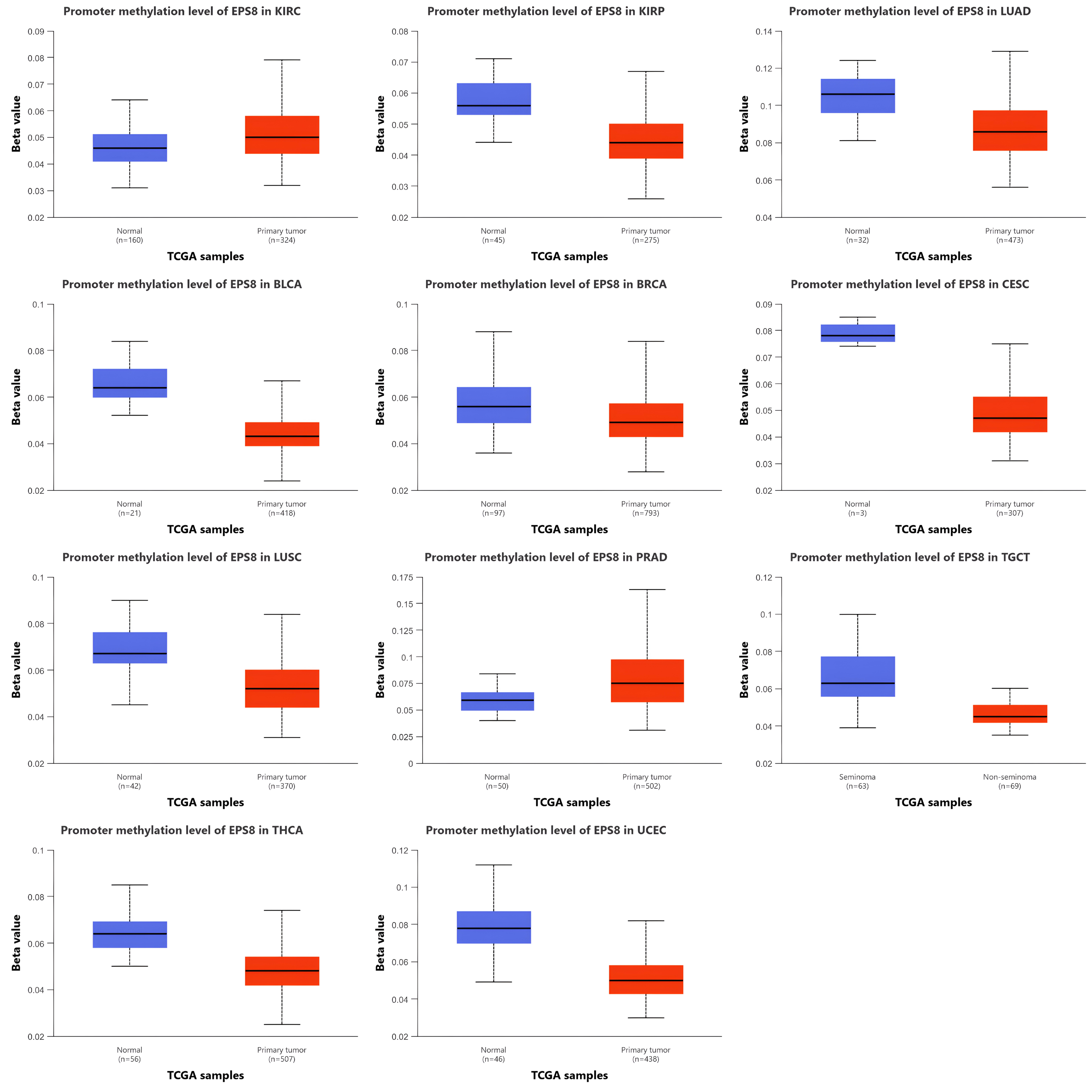

**Figure 2:** DNA methylation analysis of EPS8 analyzed using UALCAN. Promoter Methylation level was significantly upregulated in KIRC and PRAD, while significantly downregulated in BLCA, BRCA, CESC, KIRP, LUAD, LUSC, TGCT, THCA, and UCEC.

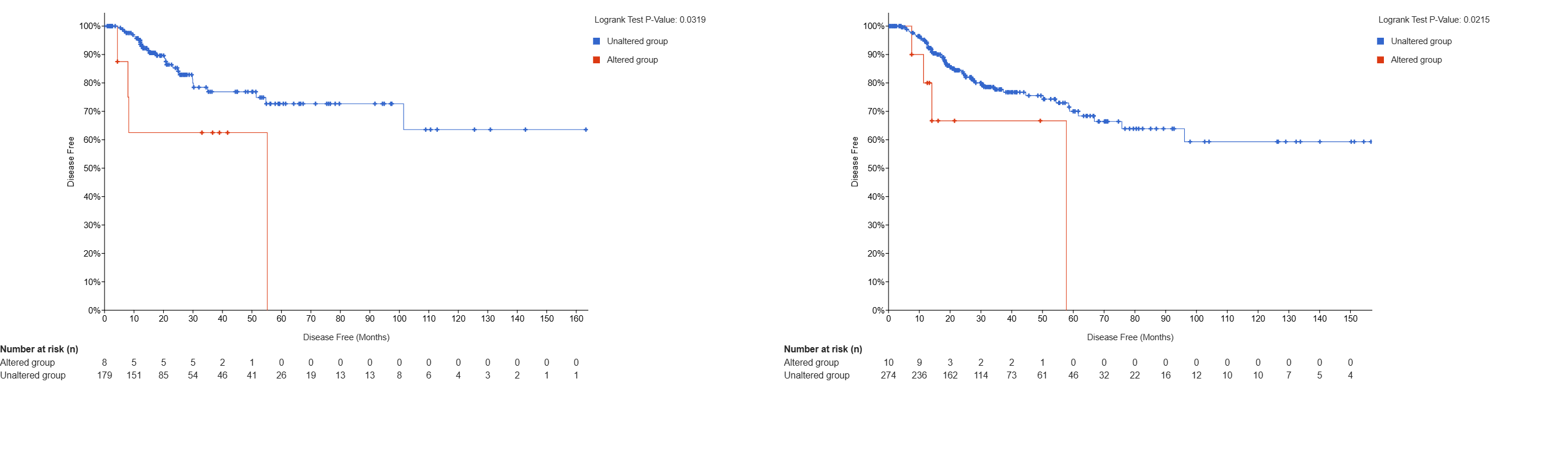

**Figure 3:** Visualization of the genetic alteration analysis of EPS8 shows poor prognosis in LUSC and BLCA patients in terms of disease-free survival.

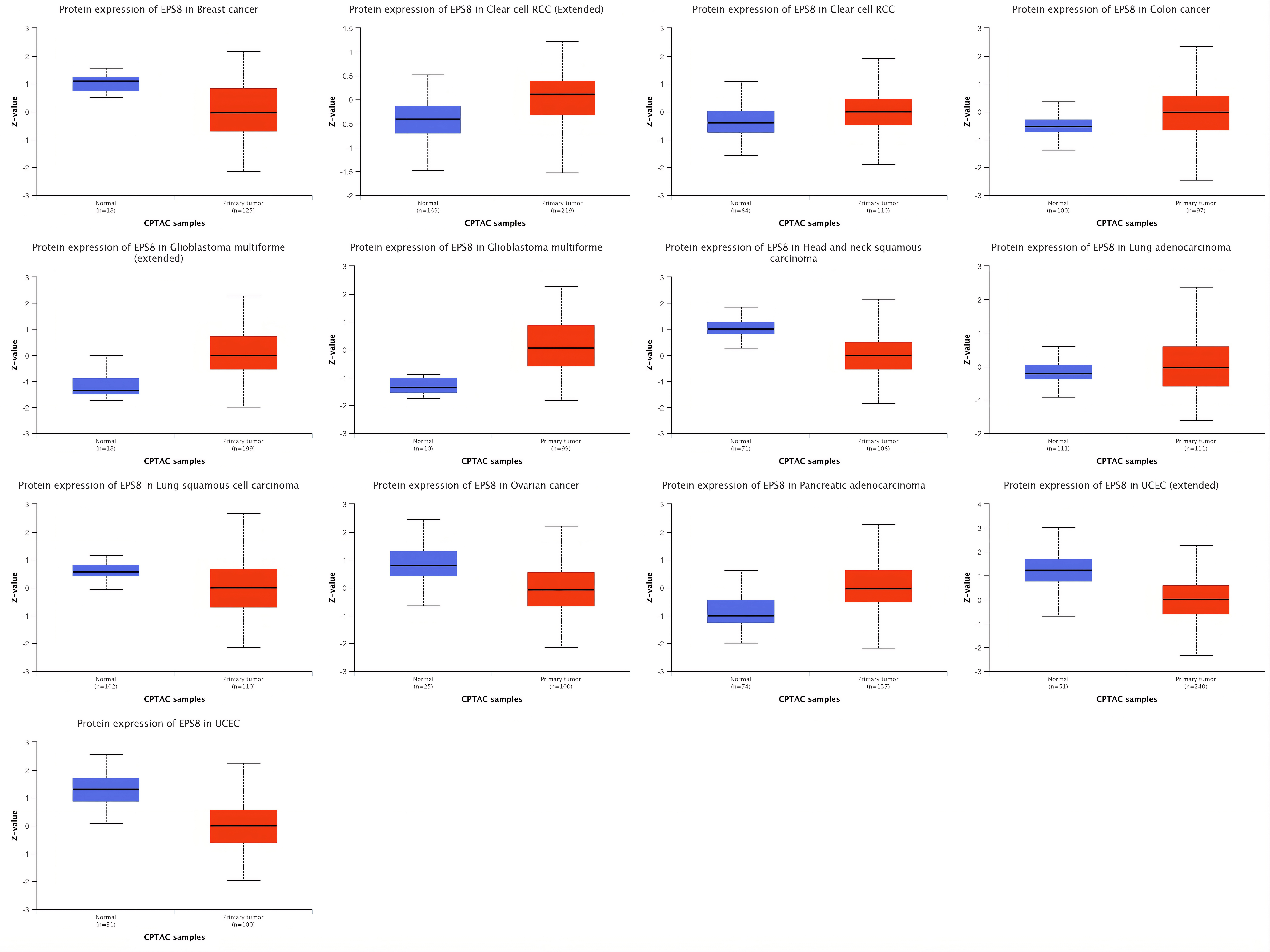

**Figure 4:** Proteomic expression analysis of BRCA, KIRC (extended), KIRC, COAD, GBM (extended), GBM, HNSC, LUAD, LUSC, OV, PAAD, UCEC (extended), and UCEC was performed using the UALCAN (https://ualcan.path.uab.edu/) database (p < 0.05).

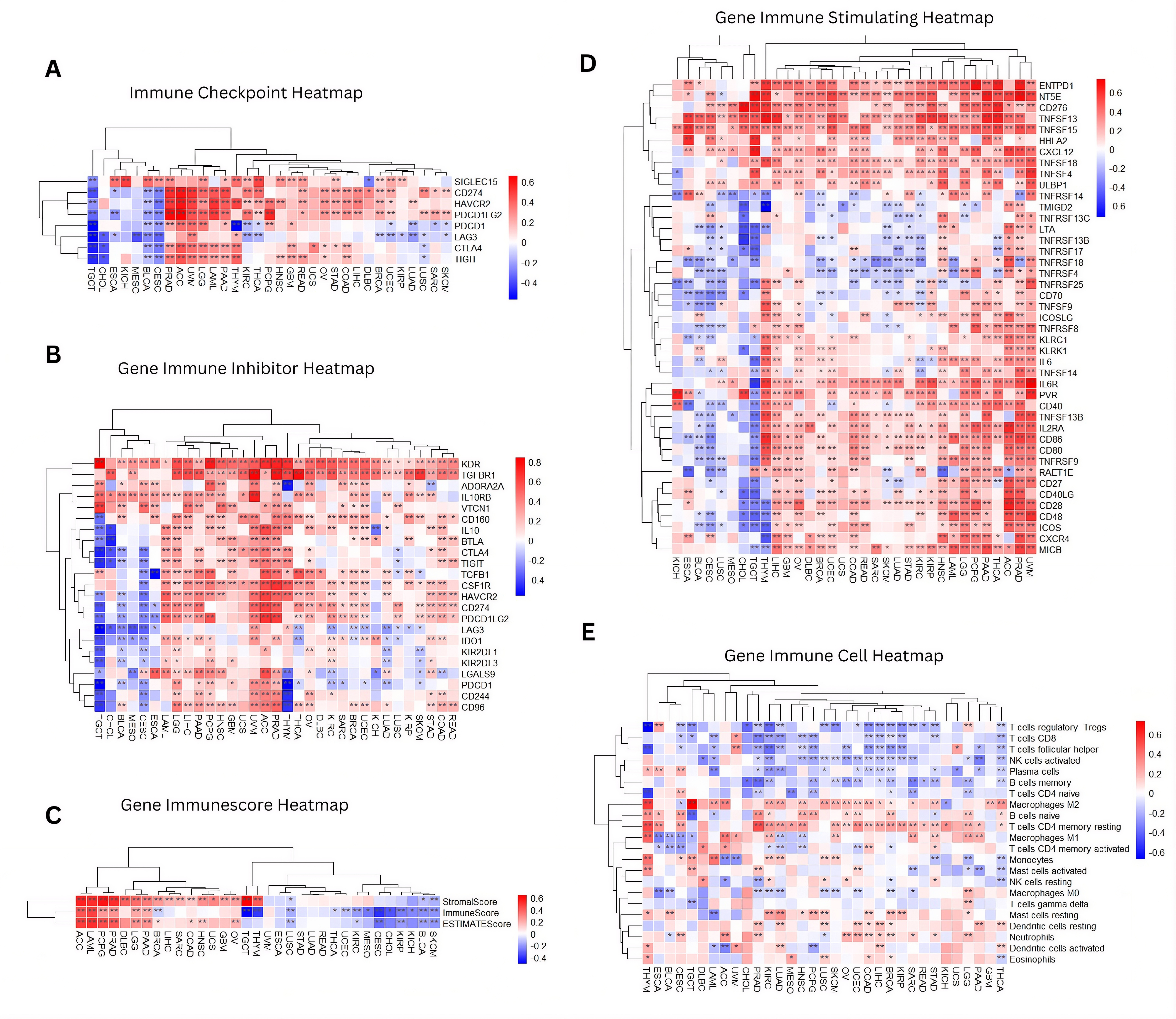

**Figure 5:** EPS8 and immune microenvironment pan-cancer correlation (A) Immune checkpoint genes heatmap, (B) Immune inhibitory genes heatmap, (C) Immune score heatmap, (D) Immune stimulating genes heatmap, (E) Immune cell Heatmap
