## Supplementary figures and images for "Comprehensive Pan-Cancer Analysis Reveals the Prognostic, Molecular, and Immunological Significance of EPS8"

### Supplement_File_Figure_01

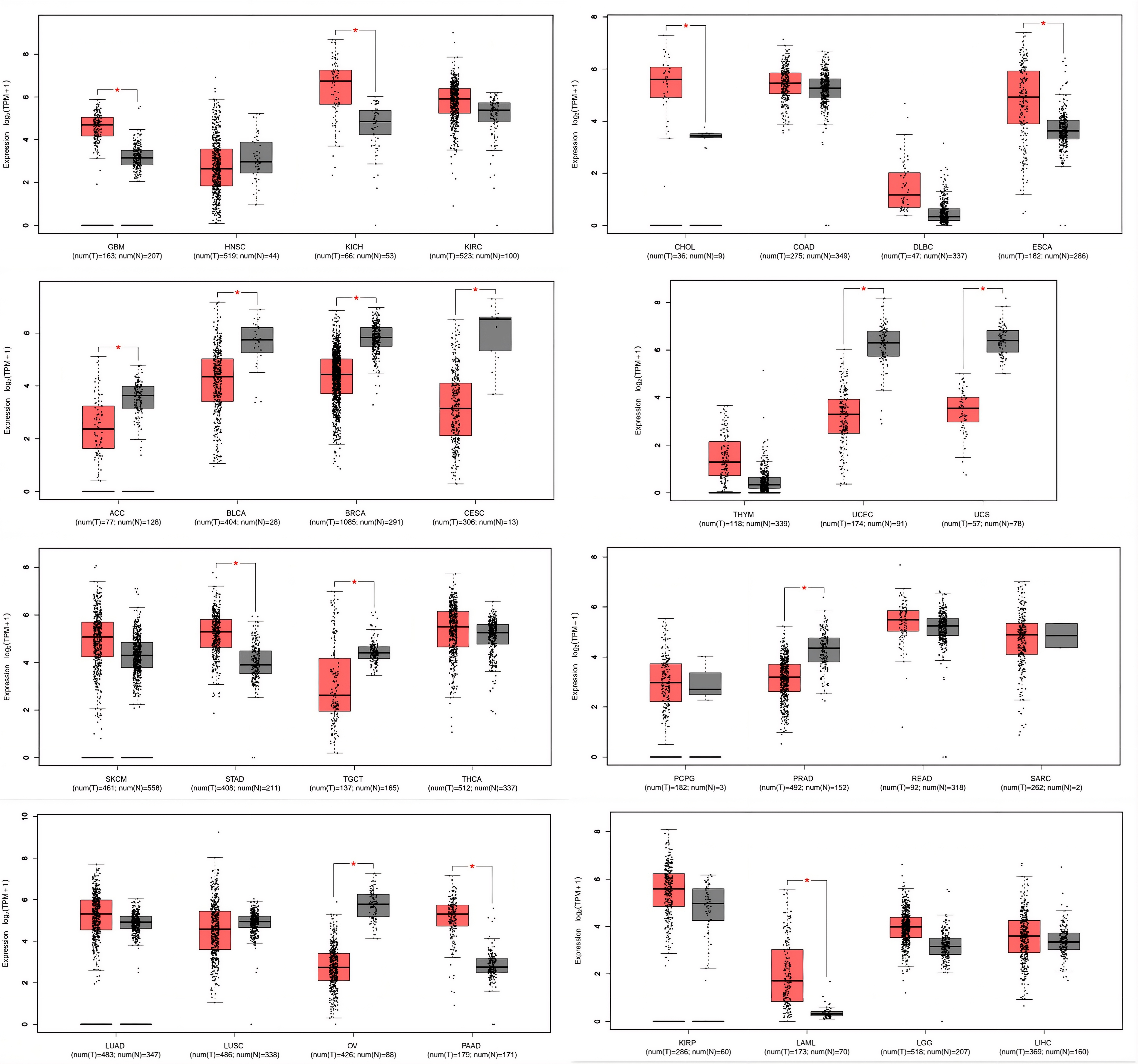

### Supplement_File_Figure_02

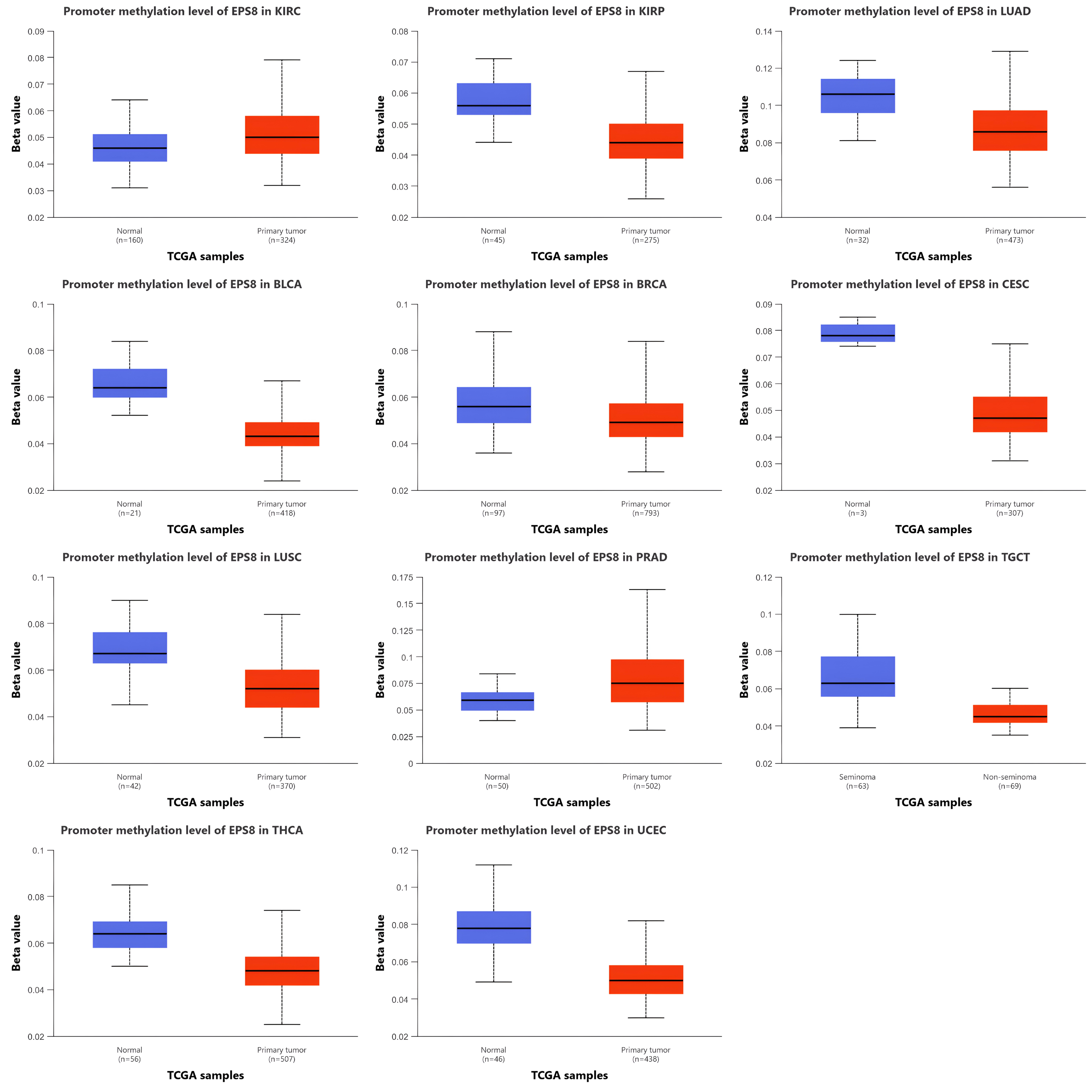

### Supplement_File_Figure_03

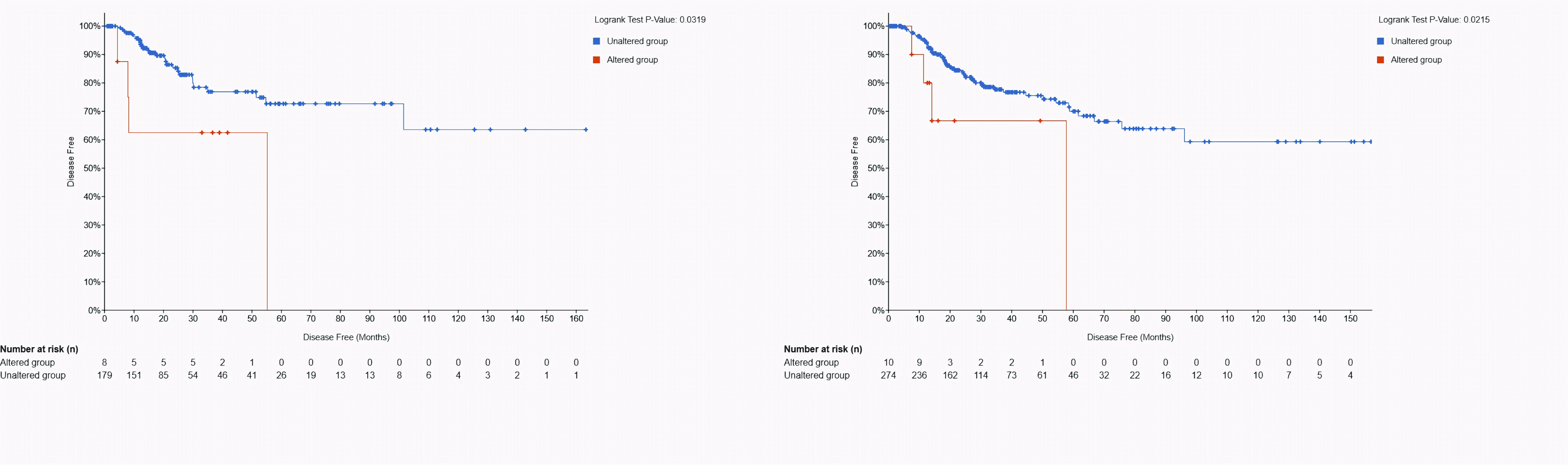

### Supplement_File_Figure_04

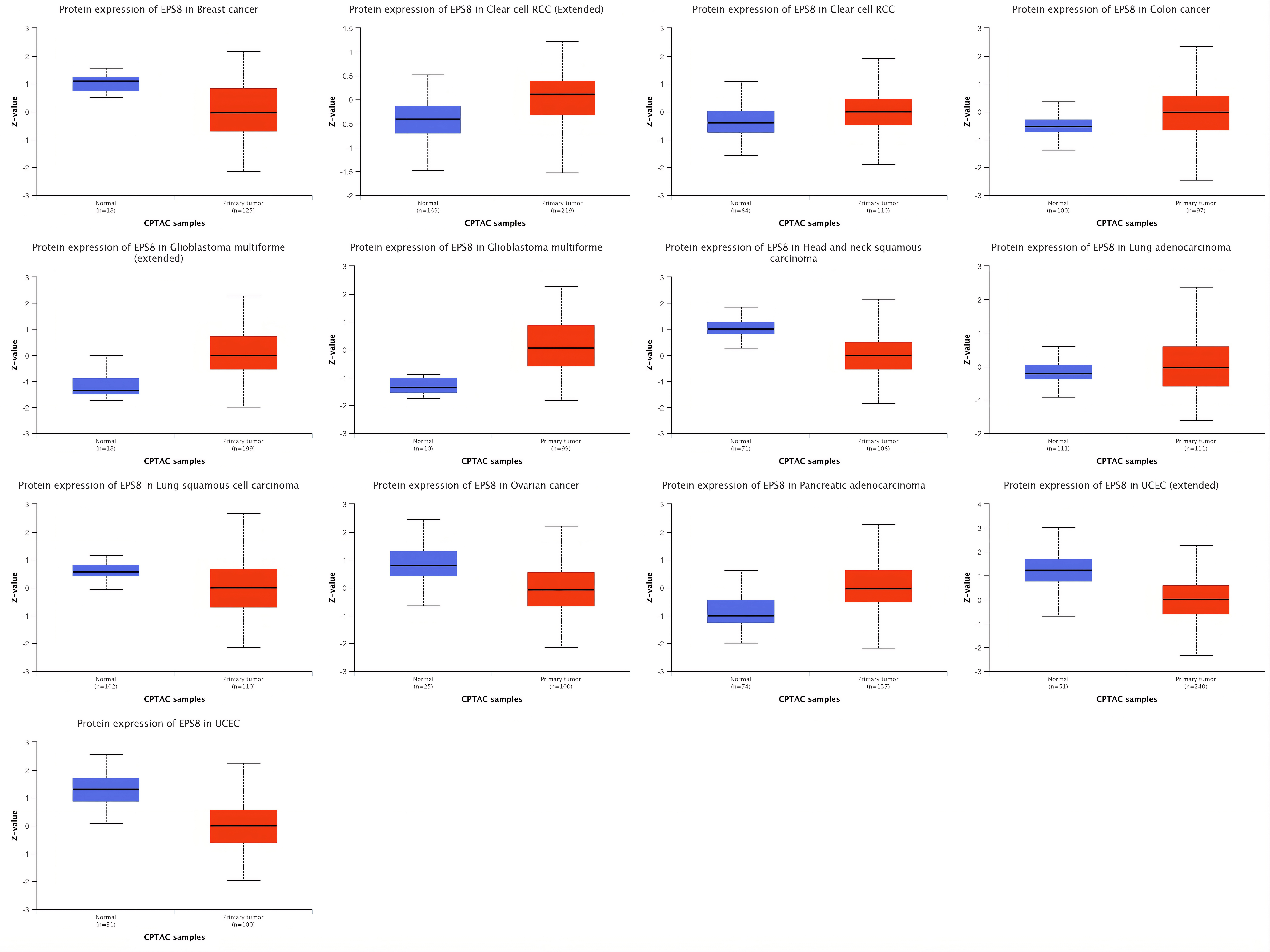

### Supplement_File_Figure_05

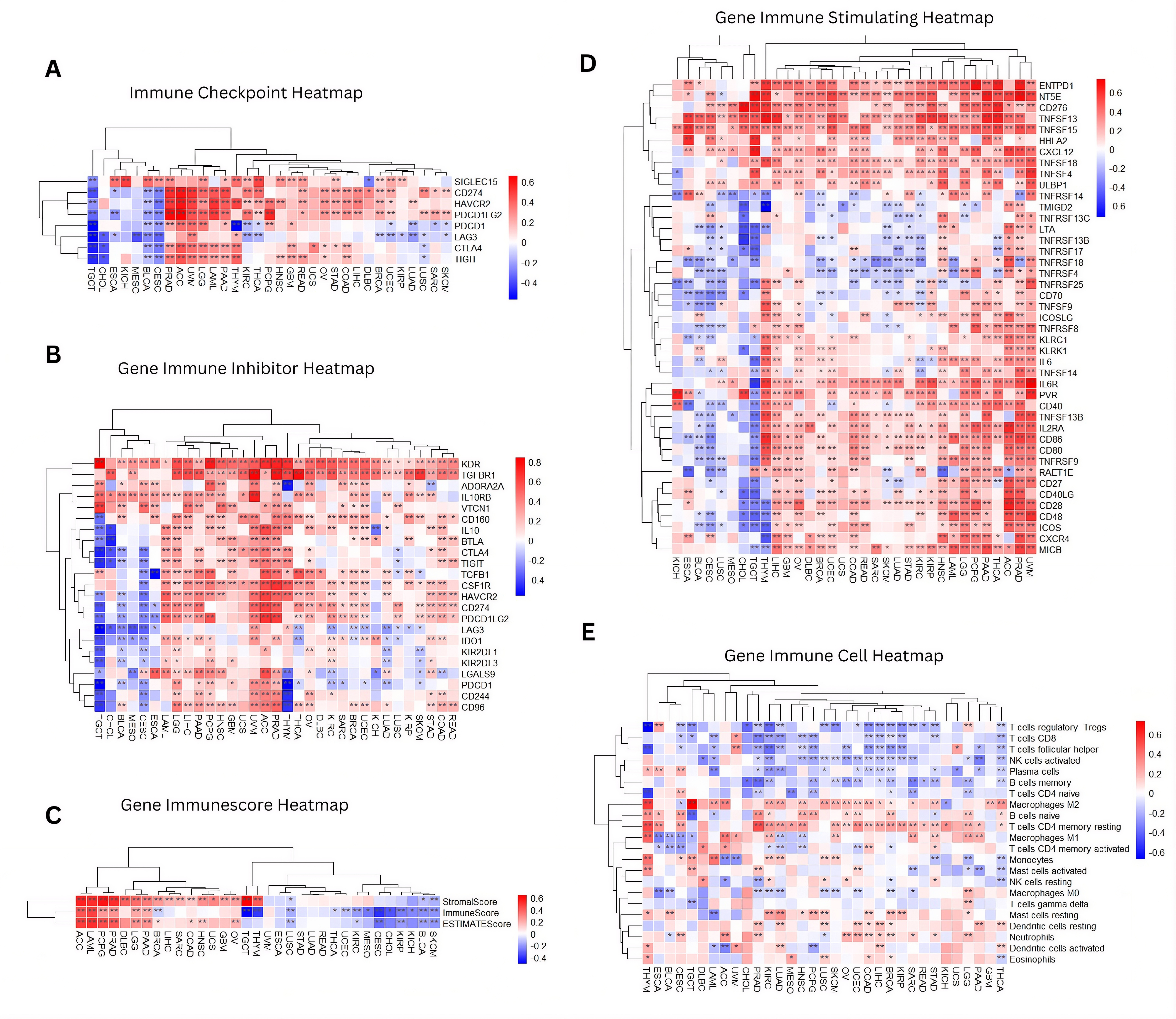
